# Acinar-to-ductal metaplasia in the pancreas requires a glycolytic switch and functional mitochondria

**DOI:** 10.1101/2022.06.27.495427

**Authors:** Thorsten Neuß, Nils Wirges, Min-Chun Chen, Sinem Usluer, Rupert Oellinger, Svenja Lier, Michael Dudek, Tobias Madl, Martin Jastroch, Katja Steiger, Werner Schmitz, Henrik Einwächter, Roland M. Schmid

## Abstract

Reprogramming of the cellular metabolism is a hallmark of pancreatic cancer, yet it remains unclear at what stage during carcinogenesis it occurs. Here, we investigated the metabolic requirements for acinar-to-ductal metaplasia (ADM), the first step in pancreatic carcinogenesis. We detected increased glycolytic marker expression in human ADM suggesting that a metabolic switch occurs during ADM formation. We report that this switch was similarly required for ADM formation in different oncogenic mouse models (KRAS, PI3K, and MEK1) and in ligand-induced ADM in mouse wild-type acini.

In addition, we show that a functional electron transport chain (ETC), but not mitochondrial ATP production, was essential to ADM formation. We conclude that the ETC provides NAD^+^ for the *de novo* synthesis of serine from glycolysis intermediates. Our findings demonstrate that metabolic programming is essential for the initiation of pancreatic carcinogenesis and thus identifies potential targets for metabolic intervention.

## Introduction

Metabolic requirements change throughout cancer progression^1^. Tumour development depends on nutrient uptake and biosynthesis. Compared with anaerobic glycolysis, oxidative phosphorylation (OxPhos) is more efficient in producing ATP. However, Otto Warburg described that cancer cells reprogram their metabolism to enhance glycolysis even under aerobic conditions, termed as aerobic glycolysis^2^. In pancreatic cancer oncogenic *Kras* favours glycolysis and pathways deriving from glycolysis. These changes provide cells with building blocks for anabolism and redox equivalents^3^. However, recent data suggest, that most cancer cells do not fully rely on glycolysis. Instead, they preserve metabolic plasticity to enable cancer cells to switch between aerobic glycolysis and OxPhos during tumourigenesis^4^. Cancer stem cells depend on OxPhos while the bulk of highly proliferative cancer favours glycolysis^5^. These concepts apply only to fully developed tumours. The metabolic state of early lesions remains unclear.

Pancreatic ductal adenocarcinoma (PDAC) is one of the most aggressive cancers and mostly incurable when diagnosed. The main reason for the dismal survival rate is that the patients become symptomatic at an advanced stage leaving only limited treatment options. Owing to the low incidence of pancreatic cancer, a population-based screening is not feasible.

However, surveillance programs have been proposed for patients with hereditary tumour predisposition. Contrary to other tumour entities, prophylactic pancreatectomy is not indicated due to its high risk of morbidity and even mortality. Hence, novel strategies for PDAC prevention need to be developed, particularly for at-risk individuals. PDAC development and progression is a multi-step process characterized by activating *KRAS* mutations in more than 90% of ductal pancreatic adenocarcinoma, sequentially followed by the inactivation of tumour suppressor genes^6^. Previously, we have shown that TGFα can induce transdifferentiation of acinar cells into duct-like cells. This acinar-to-ductal metaplasia (ADM) was proposed as the first step in pancreatic carcinogenesis^7^. Indeed, lineage tracing experiments demonstrated a mainly acinar origin of PDAC^8 9,10^. As such, acinar cells represent the origin of pancreatic intraepithelial neoplasia (PanINs). However, PDAC can in principle also originate from duct cells.

Here, we investigated the metabolic requirements for ADM. We discovered that a metabolic switch with increased glycolysis and an ATP-independent ETC activity is required for acinar-to-ductal metaplasia.

## Results

### ADM *in vivo* is accompanied by metabolic changes

The reprogramming of the metabolism to enhance glycolysis including overexpression of glycolytic enzymes is a feature which has already been described for pancreatic cancer^11^. The onset of these changes during pancreatic cancer development was investigated by analysing the expression level of glycolytic markers in human tissue microarrays containing normal tissue and ADM (Figures 1A-F). We detected significantly enhanced expression of glucose transporter type 1 (GLUT1), hexokinase 1 (HK1) and lactate dehydrogenase A (LDHA) in ADM lesions compared to normal tissue (Figures 1G-I). This increased expression suggests that the glycolytic switch already occurs during ADM formation.

**Figure 1.**
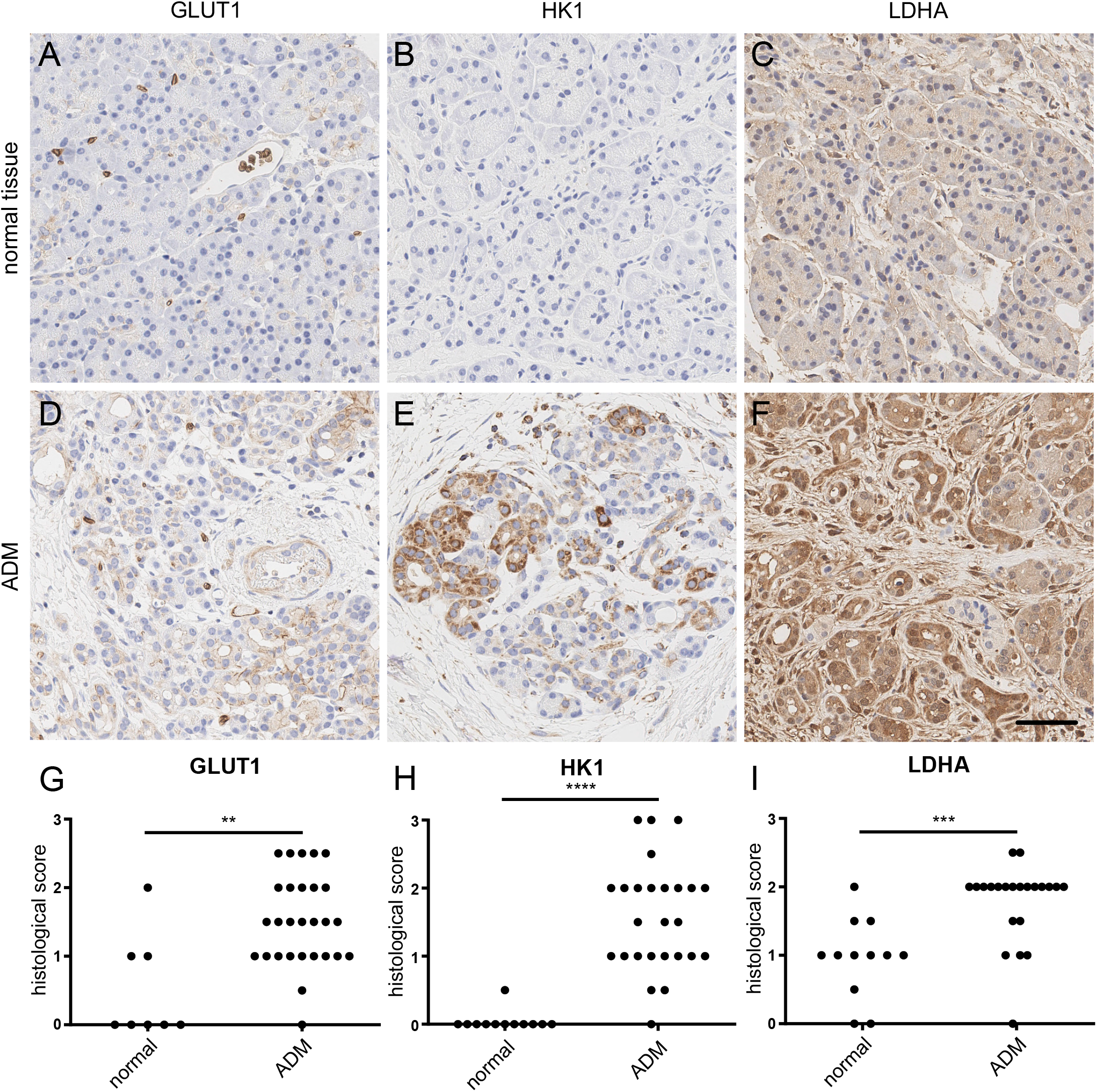
Glycolytic markers are upregulated in human ADM lesions. **A, D**. Immunohistochemistry for GLUT1, **B, E** for HK1 and **C, F** for LDHA on human tissue microarrays. Scale bar 50 μm. **G**. Histological scores for GLUT1 IHCs in A and D. **H**. Histological scores for HK1 IHCs in B and E. **I**. Histological scores for LDHA IHCs in C and F. Histological scores were given separately for normal tissue, early and late ADM lesions. *p<0.05, **p<0.01, ****p<0.0001.

To investigate whether metabolic reprogramming was common among ADM lesions, we studied three preclinical mouse models of PDAC that recapitulate human pancreatic carcinogenesis by their driver mutations as well as by their morphology. Mice that endogenously express *Kras*^*G12D*^ using *Ptf1a*^*Cre/+*^ develop ADM and present the full spectrum of PanIN lesions^12^. In addition, a gain of function mutation of *PIK3CA* (*Pi3k*^*CAH1047R*^), which occurs in cases with wild-type *KRAS*, and a gain of function mutation of *MEK1* (*Mek1dd*) results in the development of ADM and PanIN^13^. Mice with all three genotypes exhibited ADM formation at 10 weeks of age (Figures 2A-D). Quantification of these lesions revealed that compared with wild-type mice, *Kras*^*G12D*^ mice developed only a few ADM lesions, whereas *Pi3k*^*CAH1047R*^ or *Mek1dd* mice developed ADM at higher rates (3x higher in *Pi3k*^*CAH1047R*^ and 5x higher in *Mek1dd* mice, compared to *Kras*^*G12D*^) (Figure 2E). These ADM lesions stained positive for the glycolytic markers GLUT1, HK1 and LDHA (Figures 2F-G and S1A-C, E-G, I-K, M-O), which were significantly upregulated compared to normal pancreatic acinar cells of the same animal (Figures 2H-J). Notably, LDHA was already elevated in tissues from oncogene-harbouring mice that had a comparable appearance to those from wild-type animals. In contrast, the expression of ATP synthase F1 subunit delta (ATP5D), a marker of oxidative phosphorylation, did not change in ADM lesions (Figures 2K and S1D, H, L, P). These data demonstrate that glycolytic markers are induced in ADM lesions of the three murine genetic pancreatic cancer models.

**Figure 2.**
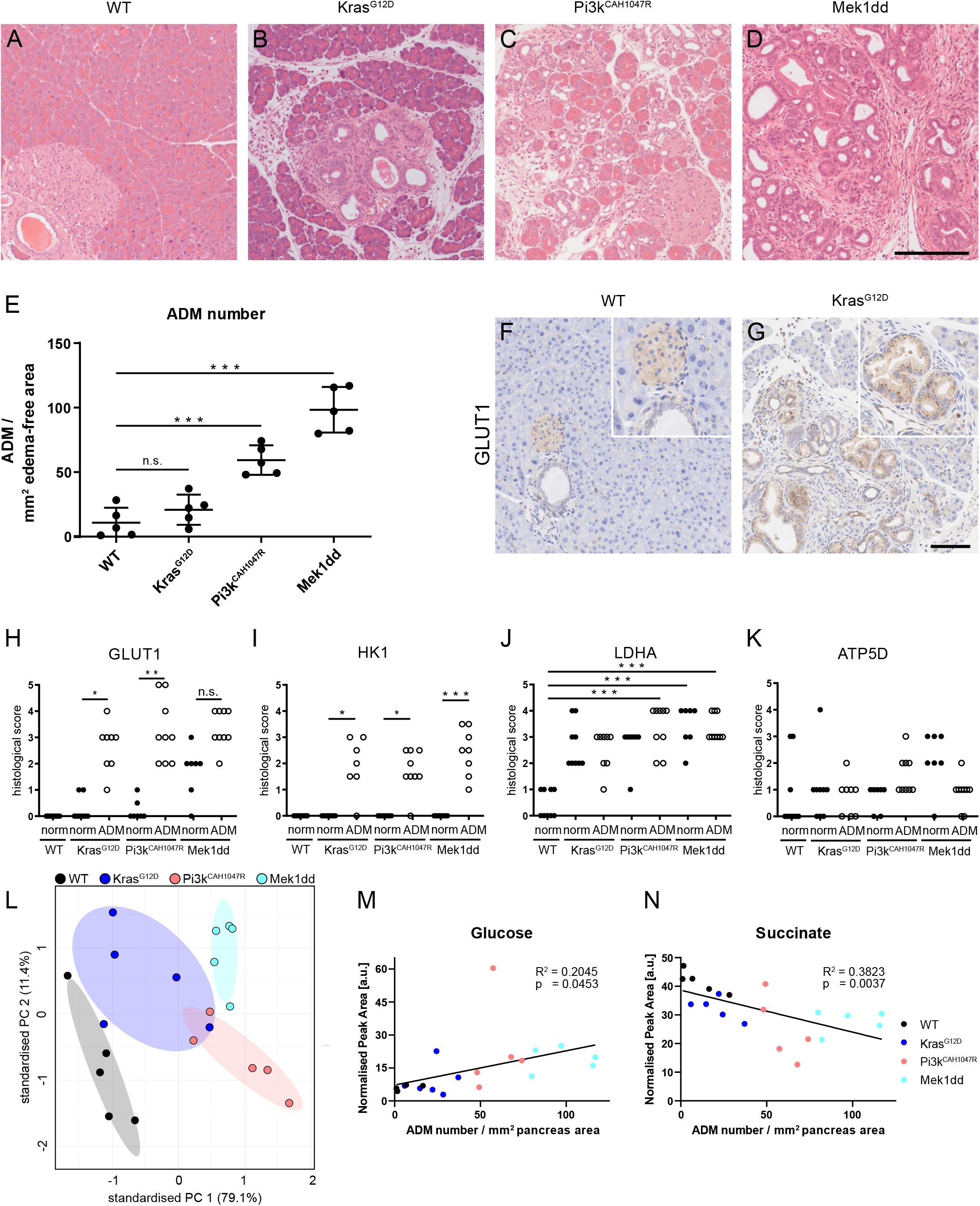
ADM formation *in vivo* is accompanied by metabolic changes. **A-D**. H&E stainings of pancreata of 10 week old animals. Scale bar = 100 μm. **E**. Number of ADM lesions per mm^2^ acinar tissue of pancreata of 10 week old mice. **F, G**. Immunohistochemistry for GLUT1 on pancreatic tissue from 10 week old mice. Scale bar = 100 μm **H-K**. Histological scores for IHCs for GLUT1 (**H**), HK1 (**I**), LDHA (**J**) and ATP5D (**K**) on pancreatic tissue from 10 week-old mice. Histological scores were given separately for normal tissue and ADM lesions. **L**. Principal component analysis of detected metabolites in tissue samples from 10 week-old animals. **M, N**. Amounts of glucose (**M**) and succinate (**N**) in the pancreas of 10 week-old animals. Metabolite amounts were plotted against the number of ADM lesions per mm^2^ of pancreatic acinar tissue. Results are mean +/- SD, *p<0.05, **p<0.01, ***p<0.001.

To analyse metabolic changes in detail, we performed nuclear magnetic resonance (NMR) metabolome analysis of tissue from 10-week-old mice of all three mouse models and wild-type animals. The principal component analysis revealed that samples clustered according to genotypes (Figure 2L), with samples exhibiting higher ADM rates showing greater differences to wild-type samples. The glucose levels in the tissue increased as the number of ADM lesions increased (Figure 2M), whereas, the level of the TCA metabolite succinate decreased (Figure 2N). Collectively, these findings indicate that a metabolic switch towards increased glycolysis is associated with ADM formation in human and murine tissue.

### Glycolysis is required for ADM formation

To further investigate these *in vivo* findings we used an *in vitro* ADM formation assay. In this well-established assay, normal acinar cells are isolated from the pancreas, embedded into a 3D collagen gel and transdifferentiated into ADM structures. In wild-type acini, ADM can be induced by the inflammatory ligands TGFα and interleukin 17a (IL17a)^7,14^, whereas, explanted acini from *Kras*^*G12D*^, *Pi3k*^*CAH1047R*^, or *Mek1dd* mice undergo ADM without further treatment (Figure S2A-E). Low ADM rates were found in ligand induced ADM. The highest transdifferentiation rates were detected in *Mek1dd*, followed by *Pi3k*^*CAH1047R*^ and *Kras*^*G12D*^ acini (Figure 3A).

**Figure 3.**
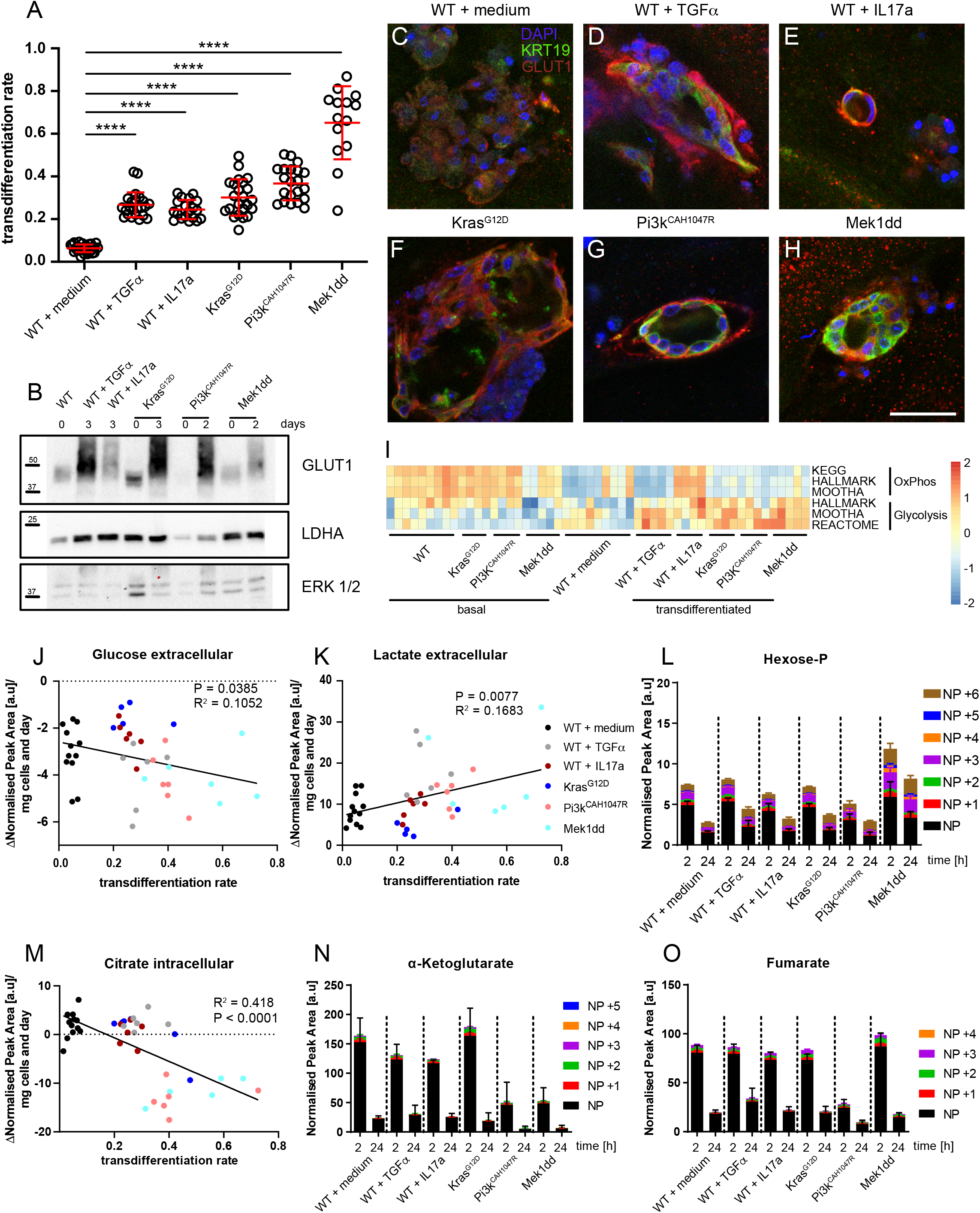
Energy metabolism changes from OxPhos to glycolysis during ADM *in vitro*. **A**. Transdifferentiation rates of wild-type acinar cells with or without stimulus and acinar cells harbouring active oncogenes. **B**. Protein expression of GLUT1 and LDHA in either freshly isolated or transdifferentiated acinar cells. ERK1/2 served as loading control. **C-H**. Immunofluorescence staining of wild-type and transdifferentiated acinar cells. Cells were stained for GLUT1 (red), KRT19 (green) and DAPI (blue). Scale bar = 50 μm. **I**. GSVA scores for genesets representing glycolysis and OxPhos in RNAseq data from freshly isolated (“basal”) and cultured acinar cells (“transdifferentiated”). **J, K**. Amounts of glucose (**J**) and lactate (**K**) in supernatants of transdifferentiated acinar cells. Values of unused medium were subtracted from the respective samples. Medium values were normalised to the pellet weight of the acinar cells. **L**. Isotopologue distribution of hexose-P 2 or 24 h after isolation of acinar cells. **M**. Amounts of citrate in transdifferentiated acinar cells. Values of freshly isolated acinar cells were subtracted from the respective samples. **N, O**. Isotopologue distribution of a-ketoglutarate (**N**) and fumarate (**O**) 2 or 24 h after isolation of acinar cells. NP = normal pattern. Results are mean +/- SD, ****p<0.0001.

The phenotypic switch of ADM is characterized by reduced expression of acinar markers (e.g. AMY2A) and enhanced expression of ductal markers such as KTR19^15^. Indeed, ductal-like structures were negative for AMY2A and stained positive for KRT19, as confirmed on RNA and protein level as well (Figures S2F-M). Downregulation of acinar genes and induction of ductal genes was also confirmed when comparing transcriptomes of freshly isolated (basal) and transdifferentiated acini (Figure S2N).

In addition to the changes in acinar and ductal genes following ADM formation, we observed increased *Glut1* expression (Figure S3A) and increased GLUT1 and LDHA protein levels in comparison to freshly isolated acini (“0 days”) (Figure 3B). Immunofluorescence staining showed that GLUT1 expression colocalised with KRT19, corroborating our *in vivo* findings of increased GLUT1 expression in ADM lesions (Figures 3C-H). A further analysis of the transcriptional changes revealed that enrichment scores of gene sets related to glycolysis were upregulated during transdifferentiation in comparison to freshly isolated acini (“basal”); whereas OxPhos-related sets were predominantly downregulated (Figure 3I). These results suggest that transdifferenting acini switch to glycolysis.

Additionally, the metabolome was analysed by NMR: Here, we observed decreased glucose levels and increased lactate levels in transdifferentiated acinar cells and their supernatants compared to freshly isolated acini. These changes correlated with the different ADM rates (Figures 3J-K and S3B-C). This finding is in line with ADM-induced glucose consumption and lactate secretion.

Isotopic tracing following the metabolism of ^13^C-labeled glucose by LC-MS analysis revealed a decrease in intracellular hexose-P after 24 h (Figure 3L). The labelling was comparable to the basal timepoint (“2 h”), indicating the consumption of glucose. In addition, TCA cycle intermediates citrate, succinate and fumarate decreased following ADM (Figure 3M, S3D-E). Isotopic tracing analysis revealed that TCA cycle metabolites like alpha-ketoglutarate and fumarate decreased in both amount and labelling (Figures 3N-O). Therefore, decreased levels TCA cycle intermediates are likely the result of reduced synthesis rather than increased consumption and turnover.

To directly assess glycolysis and mitochondrial respiration during ADM, the extracellular acidification rate (ECAR) and the oxygen consumption rate (OCR) were measured. Basal OCR was increased significantly and close to maximal rates in *Mek1dd* acini compared to all other genotypes (Figure S4A, E). In addition, ECAR was elevated at basal levels in *Mek1dd* and *Pi3k*^*CAH1047R*^ acini, suggesting an increased energy demand. Upon addition of oligomycin to block mitochondrial ATP production, ECAR increased significantly, demonstrating a pronounced spare glycolytic capacity in *Pi3k*^*CAH1047R*^ and *Mek1dd* acini (Figure 4A, S4F). ATP production rates calculated from flux measurements revealed that *Mek1dd* acini displayed highest levels of ATP production compared to all other genotypes Figure 4B). In all genotypes, glycolysis-derived ATP increased compared to medium-treated wild-type acini. The glycolysis-derived ATP followed the same trend as the transdifferentiation rates measured 24 h later supporting the association between glycolysis and ADM formation.

**Figure 4.**
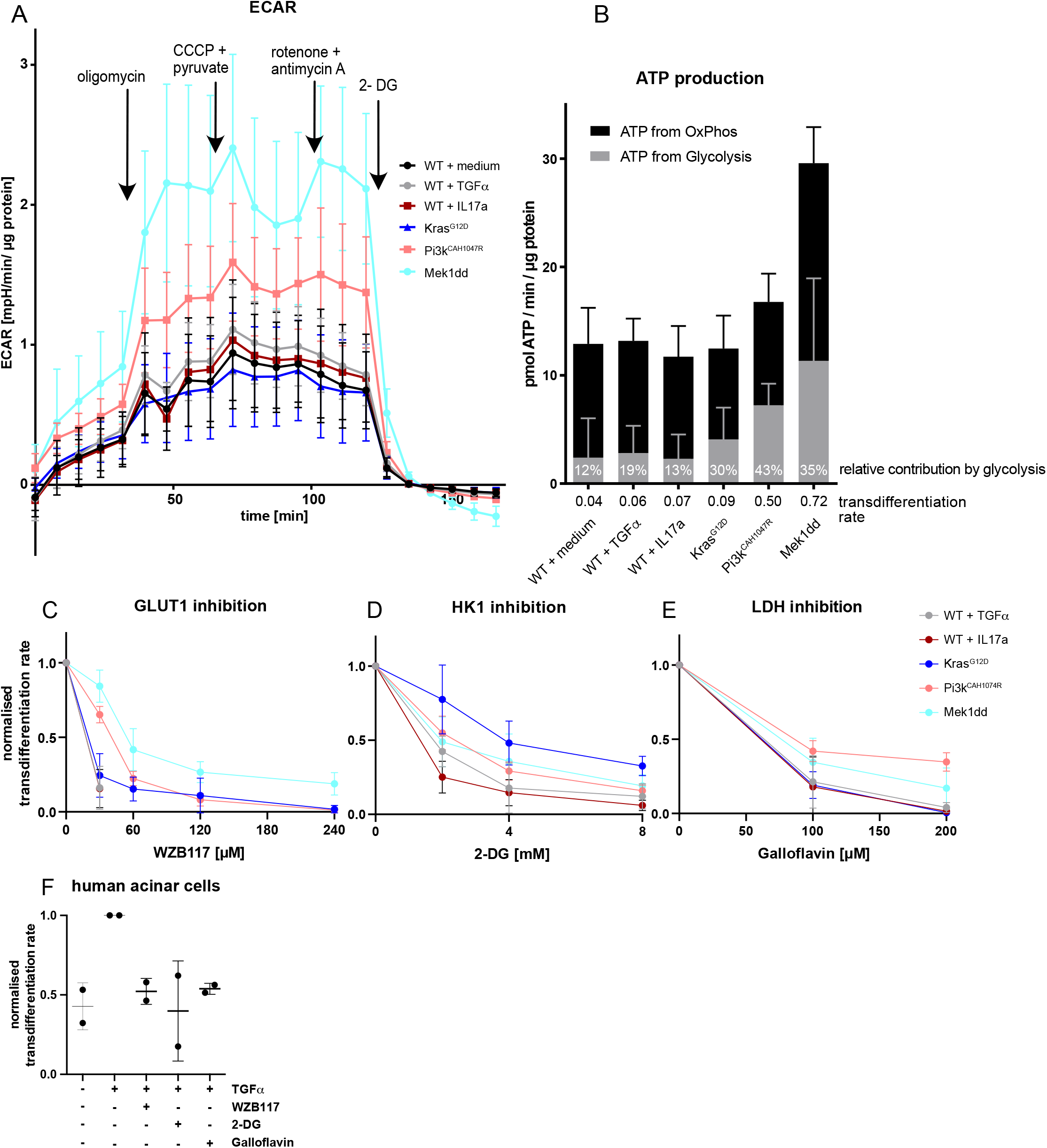
Glycolysis is essential for ADM formation. **A**. Extracellular acidification rates (ECAR) of acinar cells 24 h after isolation. Cells were treated with the displayed inhibitors at indicated timepoints. All values were normalized to protein content per well (N = 7-9). **B**. Calculated ATP production from ECAR and OCR rates in acinar cells 24 h after isolation. Contribution of ATP from glycolysis is given in percent of the total ATP production. The displayed transdifferentiation rate was assed 48 h after acinar cell isolation (N = 7-9). **C, D, E**. Transdifferentiation rates after treatment with the GLUT1 inhibitor WZB117 (N = 4-12) (**C**), HK1 inhibitor 2-deoxyglucose (2-DG) (N = 4-5) (**D**) or LDHA inhibitor galloflavin (N = 4-5) (**E**). **F**. Transdifferentiation rates of human acinar cells after stimulation with TGFα and inhibition with either WZB117, 2-DG or Galloflavin. Results are mean +/- SD.

To investigate the functional need for glycolysis during ADM, we blocked glycolysis at several levels. The GLUT1 inhibitor WZB117 effectively inhibited transdifferentiation (Figure 4C). Low transdifferentiation rates were already effectively inhibited at low WZB117 concentrations. However, higher doses of WZB117 were required to block ADM in *Mek1dd* or *Pi3k*^*CAH1047R*^ acini. Similarly, HK1 inhibition by 2-deoxy-d-glucose (2-DG) led to a dose-dependent reduction of ADM (Figure 4D) and treatment with galloflavin, an LDHA inhibitor, reduced ADM as well (Figure 4E). We observed no treatment-induced cytotoxicity in acini treated with glycolysis inhibitors (Figure S4B-D). To validate the murine findings, human acini were isolated from healthy pancreata and exposed to TGFα (Figure 4F). Similar to murine acini, WZB117, 2-DG or galloflavin inhibited ADM formation in human acini.

Taken together, acinar cells undergoing ADM increase their glycolytic capacity *in vitro* and *in vivo*. This metabolic switch is a functional requirement during murine and human ADM formation.

### c-MYC reduces glucose flux to the mitochondria

In addition to the metabolic changes, we analysed expressional changes during ADM. Specifically, we looked for transcription factors which are involved in metabolic reprogramming. Strikingly, c-Myc (MYC) target genes were upregulated following ADM (Figure 5A), especially in ADM from acini harbouring oncogenes. Notably, increased MYC protein levels were detected as early as 3 h after isolation (Figure 5B). The MYC inhibitor 10058-F4 attenuated ADM formation independently of the stimulus; however, at low 10058-F4 concentrations, ADM formation in *Mek1dd* of *Pi3k*^*CAH1047R*^ acini was only slightly inhibited (Figure 5C). No changes in cell death were observed following MYC inhibition (Figure S5A). Likewise, acinar cells with a heterozygous deletion of MYC showed attenuation of ADM when exposed to TGFα or IL17a (Figure 5D). These data demonstrate that MYC is essential for ADM development.

**Figure 5.**
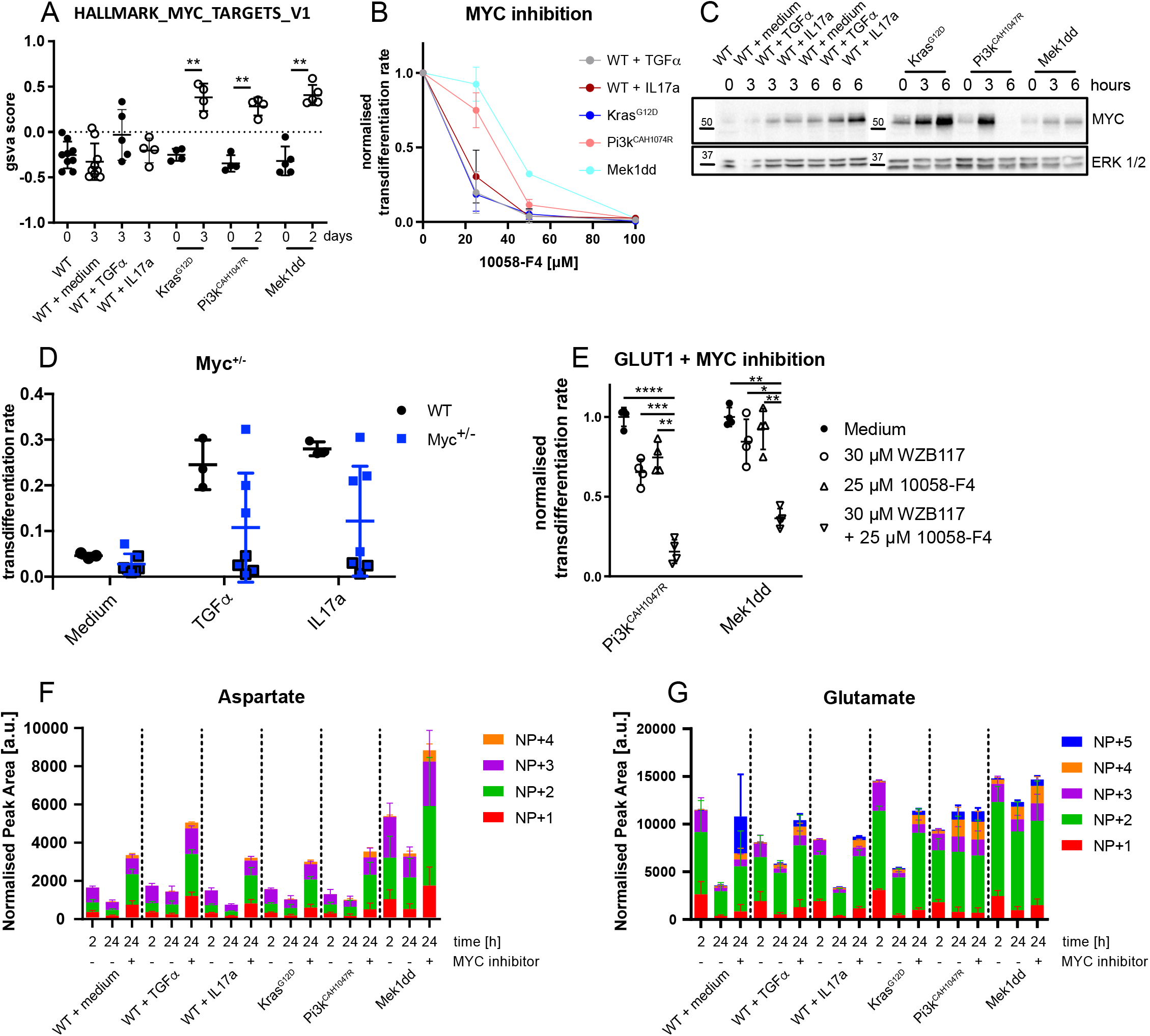
MYC regulates glucose consumption and OxPhos. **A**. Enrichment scores for the geneset Hallmark_MYC_targets_V1 in RNASeq of freshly isolated and cultured acinar cells. **B**. Western Blot for MYC of acinar cells either freshly isolated or after 3 or 6 h of incubation. ERK1/2 served as loading control. **C**. Transdifferentiation rates after treatment with the MYC-inhibitor 10058-F4 (N = 4-8). **D**. Transdifferentiation rates of wild-type and Myc^+/-^ acinar cells after induction with either TGFα or IL17a. **E**. Transdifferentiation rates of Pi3k^CAH1047R^ and Mek1dd after treatment with either GLUT1 inhibitor WZB117 or MYC inhibitor 10058-F4 or in combination. **F, G**. Isotopologue distribution of labelled aspartate (**F**) and glutamate (**G**) after treatment with ^13^C-glucose and 50 μM 10058-F4 (N = 5). NP = normal pattern. Results are mean +/- SD. **p<0.01.

To determine the effect of MYC on the glycolytic phenotype, we combined MYC and GLUT-1 inhibition during ADM formation. Whereas treatment of *Mek1dd* or *Pi3k*^*CAH1047R*^ acini with low 10058-F4 or WZB117 concentrations resulted only in minor effects on transdifferentiation (25% or 5% reduction for 10058-F4 and 35% or 15% reduction for WZB117 in PI3K^CAH1047R^ or Mek1dd, respectively), the combination of both inhibitors caused a significant synergistic attenuation of ADM (Figure 5E). The combination of both inhibitors caused no increased cell death (Figure S5B). Additionally, treatment of acini with 10058-F4 displayed a decreased in hexose-P levels as detected by ^13^C-glucose isotopic tracing (Figure S5C). Investigating the fate of the metabolised hexose-P, we found decreased labelled aspartate and glutamate after 24 h, which are both side branches of the TCA cycle (Figures 5F, G). Importantly, 10058-F4 treatment reversed this decrease indicating that MYC was involved in directing the metabolic flux away from the mitochondria during ADM.

Multiple interactions of MYC with cell metabolism have been demonstrated^16^ including not only glycolysis but also glutamine metabolism. Enzymes of the glutaminolysis like glutaminase (*Gls*), *Glud* and *Gpt2* as well as transporters for glutamine and glutamate were either unchanged or downregulated during ADM and only *Got2* was upregulated (Figure S5D). Additionally, intracellular amounts of glutamine and glutamate decreased during transdifferentiation independently of the transdifferentiation rate (Figure S5E-F). Also, extracellular levels of glutamine were independent of the transdifferentiation rate (Figure S5G-H). However, the extracellular glutamate levels increased during ADM, which was less pronounced at higher transdifferentiation rates (Figure S5H). Blocking GLS using BPTES which inhibits glutamine to glutamate conversion did not affect ADM (Figure S5I).

Taken together, the metabolic role of MYC during ADM rather appears to be the strengthening of glycolysis than induction of glutaminolysis.

### Mitochondrial activity is needed for ADM formation

So far the preserved respiration seen in the OCR measurements seems unclear considering increased glycolysis, the reduction of TCA metabolites and independence of glutaminolysis. The first step of oxidative phosphorylation (OxPhos) is performed by the pyruvate dehydrogenase (PDH). The amount of phosphorylated (inactive) PDH decreased during transdifferentiation (Figure S6A) and PDH inhibition reduced ADM formation in all stimulating conditions (Figure S6B).

Additionally, inhibition of complexes I, III, or V of the electron transport chain (ETC) using rotenone, antimycin A, or oligomycin, respectively, attenuated transdifferentiation (Figure 6A). This data clearly demonstrate that, besides glycolysis, mitochondrial function is essential for ADM formation. Surprisingly, mitochondrial uncoupling with carbonyl cyanide m-chlorophenylhydrazone (CCCP) did not affect the transdifferentiation rates (Figure 6B). Importantly, addition of CCCP to oligomycin-treated acini partially rescued the transdifferentiation of TGFα-stimulated wild-type or *Kras*^*G12D*^ acini and fully rescued ADM formation in the other inducers (Figure 6B). These data demonstrate that the ETC is required whereas mitochondrial ATP production is dispensable during ADM.

**Figure 6.**
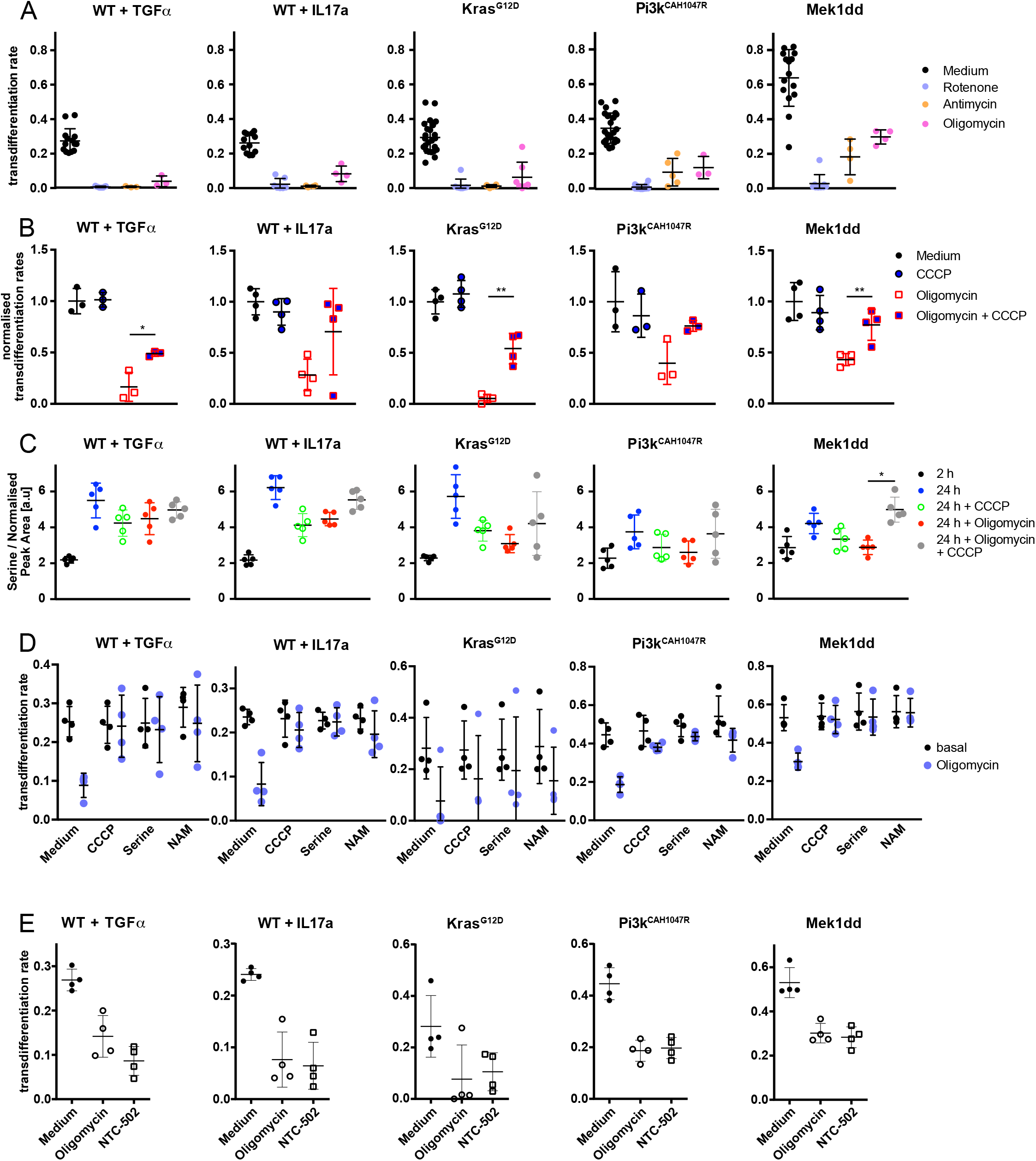
Mitochondrial metabolism, but not OxPhos is required for ADM. **A**. Transdifferentiation rates of acinar cells after treatment with ETC inhibitors rotenone, antimycin or oligomycin. **B**. Transdifferentiation rates of acinar cells after treatment with ETC inhibitors oligomycin or carbonyl cyanide m-chlorophenylhydrazone (CCCP) or in combination. Values were normalised to untreated controls. **C**. Intracellular serine amounts in acinar cells 2 or 24 h after isolation as determined by LC-MS. Acinar cells were treated either with oligomycin or CCCP or both. **D**. Transdifferentiation rates after treatment with ETC inhibitor oligomycin in combination with CCCP or serine or nicotinamide (NAM). **E**. Transdifferentiation rates after treatment with serine synthesis inhibitor NTC-502 or oligomycin. Results are mean +/- SD. *p<0.05, **p<0.01, ***p<0.001.

To understand the requirement of a functional ETC, we performed isotope tracing on acini treated with oligomycin, CCCP or both using fully labelled ^13^C-glucose. Afterwards, metabolites were screened for a pattern correlating with ADM formation: Decrease with oligomycin and rescue by CCCP. As a result, we found that intracellular serine was strongly increased during ADM (Figure 6C). This increase was partially blocked by oligomycin and rescued by CCCP paralleling the transdifferentiation rates. Isotopic tracing revealed that parts of the serine were synthesised from glucose within 2 h of labelling (Figure S6C). As a functional test, we supplemented serine in a concentration comparable to Dulbecco’s modified Eagle’s medium (DMEM) to rescue the oligomycin-induced block in ADM formation. Whereas *Kras*^*G12D*^ acini only showed a weak response, all other conditions demonstrated a partial to full rescue (Figure 6D). It has been reported already that an impairment of the ETC results in a decrease of NAD^+^ which is required for serine synthesis. To this end, we treated acini with nicotinamide (NAM), a membrane-penetrating precursor of NAD^+^. The treatment with NAM rescued the oligomycin-induced block in ADM formation, paralleling the CCCP and serine treatment. Additionally, inhibition of serine biosynthesis by NTC-502 attenuated ADM formation comparable to oligomycin (Figure 6E).

These results indicate that a functional ETC is mandatory for ADM formation by providing NAD^+^ for serine synthesis.

Serine is involved in multiple pathways like synthesis of DNA, lipids and membranes, methylation processes and ROS defence. Comparing transcriptional levels of freshly isolated and transdifferentiated acini revealed no significant upregulation of gene sets covering serine-glycine-one-carbon-metabolism (SGOC)^17^, DNA or lipid biosynthesis (Figure S6D-S6G). In contrast, genes associated with glutathione (GSH) regeneration were significantly upregulated (Figure S6H) and the precursor of GSH, cystathionine, increased in abundance as well (Figure 7B). Strikingly, both serine and cystathionine levels decreased after MYC-inhibition (Figure 7A-B), linking the sensitivity of ADM towards MYC-inhibition and serine depletion. At the same time, MYC-inhibition led to an increase in the mRNA for *Gclc*, the rate-limiting enzyme of GSH synthesis (Figure 7C). Compensatory increases of *Gclc* have been described in situations of GSH-depletion.

**Figure 7.**
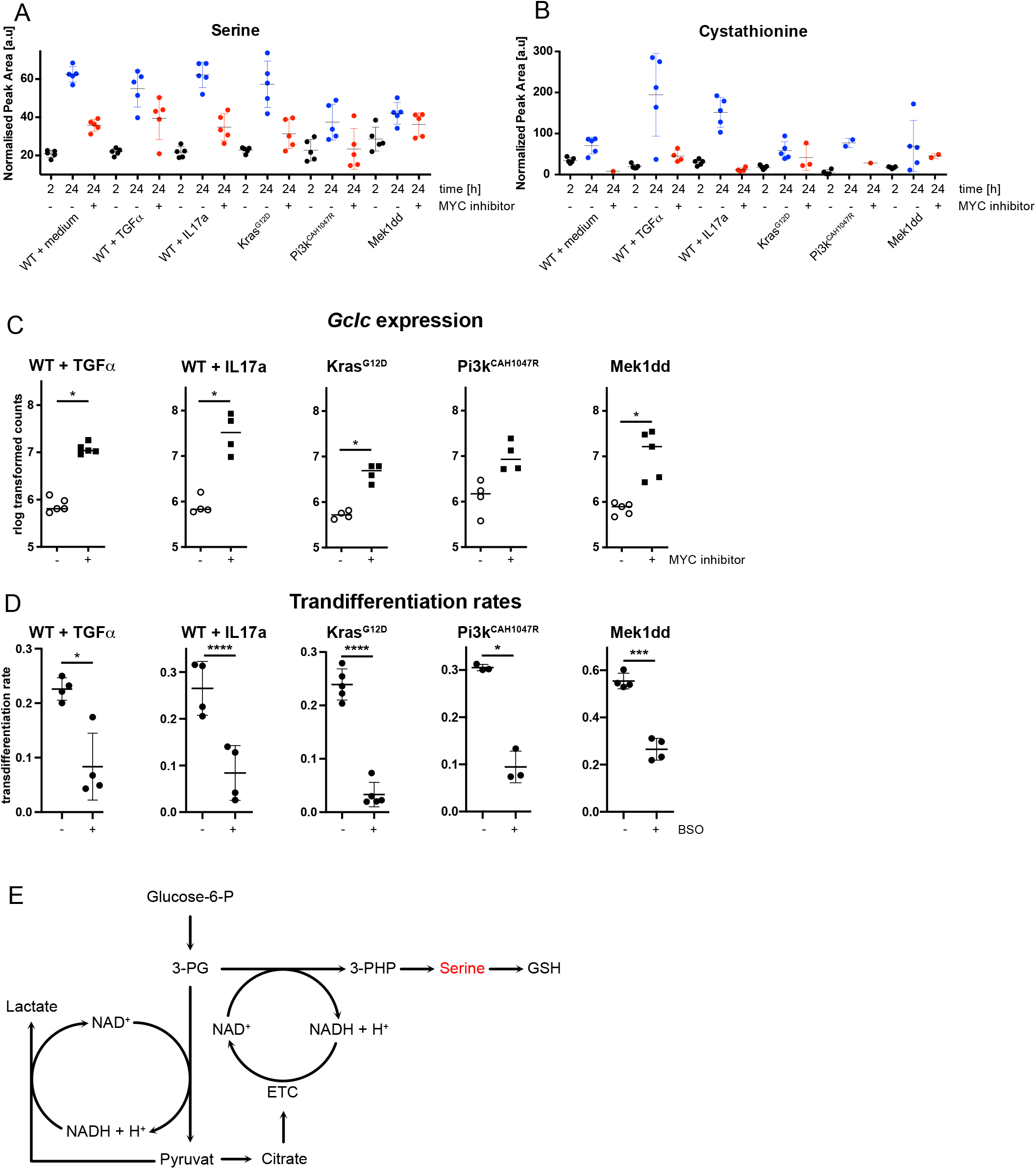
MYC-dependent *de novo* serine fuels glutathione metabolism. **A-B**. Intracellular amounts of serine (**A**) and cystathionine (**B**) in acinar cells 2 or 24 h after isolation or after MYC inhibition as determined by LC-MS. **C**. mRNA expression of the catalytic unit of glutamate-cysteine ligase (*Gclc*) in acini with and without MYC inhibition. **D**. Transdifferentiation rates after treatment with BSO. **E**. Schematic overview of serine synthesis from glucose (3-PG = 3-phosphoglycerate, GSH = glutathione). Results are mean +/- SD. *p<0.05, **p<0.01, ***p<0.001.

Additionally, we inhibited GCLC by BSO (Buthioninsulfoximin) to test for the functional relevance of GSH synthesis during ADM formation. In line with our previous data, inhibition of GSH synthesis led to a significant impairment of ADM formation in all tested conditions (Figure 7D).

These results suggest that during ADM, MYC-mediated serine synthesis is needed for GSH production.

## Discussion

One of the critical challenges of cancer prevention is the lack of understanding of key molecular changes during cancer initiation. Reprogramming of cellular metabolism is a recognized hallmark of cancer^1^. Cancer cells are dependent on these metabolic changes to maintain energy balance and support biosynthesis^18^. Unlike the abundant work on genomic and transcriptional alterations in precancerous tissues, a comprehensive analysis of metabolic reprogramming has not been performed yet.

Here, we show that murine and human ADM lesions display increased expression of glycolytic markers GLUT1, HK1, and LDHA. Increased expression of GLUT1 is invariably detected in *KRAS* mutant pancreatic lesions^19^. HK1 and 2, rate limiting enzymes in the glycolytic pathway, are upregulated in primary pancreatic cancer specimens and lactate dehydrogenase A (LDHA), executing the final step of aerobic glycolysis, is elevated in pancreatic cancer patients^20^. The increased expression of these glycolytic markers in precursor lesions of pancreatic cancer suggests that a metabolic switch, increased aerobic glycolysis and attenuated OxPhos (Warburg effect), might already occur during ADM. In addition, the amount of glucose in the tissue increased with the number of ADM lesions. At the same time, the amount of succinate, a metabolite of the TCA cycle decreased. *In vitro* studies revealed that glucose consumption and lactate secretion correlated with the transdifferentiation rate. Meanwhile, TCA metabolites decreased during ADM. Analysis of the oxygen consumption rate (OCR) and extracellular acidification rate (ECAR) revealed a greater increase in glycolysis-derived ATP with increasing ADM rates compared to OxPhos-derived ATP. Finally, glycolysis inhibition blocked ADM formation of isolated murine and human acini.

Our findings are consistent with previous observations that freshly isolated wild-type acinar cells rely on both glycolysis and OxPhos^21^, with glycolysis inhibition having a major impact on ATP homeostasis. Also, hypoxia increased glycolysis in acinar cells^22^. Moreover, cerulein-treatment, which causes hyperstimulation of acinar cell and ADM, reduced OxPhos in wild-type acini^23^.

Only a few descriptive studies indicate that cell metabolism may be reprogrammed at early stages of tumourigenesis. For example, hexokinase expression was described in liver neoplasia^24^, whereas preneoplastic rat renal basophilic cell lesions display increased Glut1 expression^25^. GLUT1 expression increases from low- to high-grade dysplasia in colon adenomas^26^. The comparison of precursor lesions and lung cancer samples indicated an early metabolic change as well^27^. A similar study was performed in hepatocellular cancer patients^28^. An increase in metabolic alterations was observed during tumour progression in these two studies. However, in both studies the “normal tissue” which was used as a control tissue was taken from resections of cancer patients. Therefore, paracrine metabolic alterations induced by neoplastic lesions in close proximity cannot be excluded. This limitation also applies to our analysis of human precursor lesions. In contrast, our murine *in vitro* data from stimulated wild-type acinar cells or active oncogenes as sole drivers exclude the influence of neighbouring tumour cells.

From these complementary results in human and mouse samples, we conclude that a metabolic switch occurs during ADM and is required for ADM formation. We propose that metabolic reprogramming is essential for the initiation of pancreatic cancer and a major driver in its progression.

Glutamine is a major source of energy entering the TCA cycle via α-ketoglutarate (α-KG). It is known that pancreatic cancer cells heavily rely on glutamine^29^. In contrast, our data demonstrate that acinar cells undergoing ADM do not depend on glutamine. Several glutamine transporters including SLC1A5 (ASCT2), SLC38A5 and the mitochondrial glutamate transporter SLC25A1 as well as enzymes of the glutamine metabolism like GLUD1, GLS and GOT are upregulated in many types of cancer^30^. Following ADM development, these transporters and enzymes were either unchanged or downregulated, with only GOT2 being slightly increased. Moreover, intracellular and extracellular amounts of glutamine did not correlate with transdifferentiation rates. Blocking the enzymatic conversion of glutamine to glutamate using the GLS inhibitor BPTES did not inhibit ADM formation. Thus, our data suggest that glutamine metabolism is not essential during ADM. It might be hypothesized that enhanced glutamine metabolism might be beneficial for cancer cells by promoting proliferation but is not required for transdifferentiation.

The transcription factor MYC is an essential downstream effector of oncogenic KRAS in the pancreas^31^. Oncogenic levels of MYC reprogram cellular metabolism, and are an integral part of the adaptation of cells to nutrient deprivation, a hallmark of cancer development. *In vivo* studies confirmed that MYC impacts the metabolism of glutamine mainly by enhancing the expression of GLS^32^. However, our data suggest that glutamine metabolism is not essential during ADM. A reason for this discrepancy might be the only transient MYC-activation observed in our system.

Recent data from PDAC cells demonstrated a MYC-dependent regulation of TCA cycle activity^33^. Additionally, upregulation of MYC favours glycolysis over OxPhos in cancer stem cells^34^. These results are in line with our finding that MYC supports glycolysis during ADM. Apart from glycolysis, serine metabolism is required for ADM formation. Serine contributes to a number of processes including protein, lipid and nucleotide synthesis, glutathione and NADPH generation and the provision of one carbon units for the folate cycle and methylation^35^.

*De novo* synthesis of serine from glucose starts by the oxidation of 3-phosphoglycerate (3-PG) which requires NAD^+^. Lactate production alone only restores as much NAD^+^ as needed for the glycolysis. Hence, additional regeneration of the NAD^+^ pool requires a functional ETC (Figure 7E). Inhibition of the ETC either pharmacologically or by hypoxia has been shown to disturb the NADH/NAD^+^ balance^36^. Therefore, *de novo* serine synthesis is dependent on a functional ETC^37^. We show that oligomycin blocks ADM formation, however, uncoupling with CCCP rescued oligomycin-inhibited ADM formation. These data indicate that ATP production is unlikely to be the main function of mitochondria during ADM. In addition, supplementing serine as well as addition of NAM was also able to rescue the inhibitory effect of oligomycin, while the effect of serine synthesis inhibition was comparable to oligomycin. Therefore, our data suggest that mitochondria are needed to provide NAD^+^ for *de novo* serine synthesis. To our knowledge, a requirement for serine synthesis in early lesions has so far not been demonstrated. In fully developed PDAC, serine was shown to be essential for tumour survival^38^. Tumours which are unable to synthesise serine are dependent on extracellular supplementation. Our culture medium did not contain serine. Therefore, our *in vitro* experiments were performed under serine-depleted conditions.

As for the fate of serine, gene sets covering DNA, lipid or membrane synthesis or one-carbon metabolism were either unchanged or downregulated. On the other hand, we demonstrate that increased intermediates of GSH during ADM formation were MYC-dependent. In addition, we could show that inhibiting GSH synthesis blocked ADM formation.

The requirement for serine synthesis in the initiation of early lesions might open strategies to prevent early pancreatic carcinogenesis with dietary of pharmacologic interventions. A serine-free diet in combination with a serine synthesis inhibitor resulted in reduced tumour growth in a *Kras*^*G12D*^-driven mouse model of colorectal cancer^39^. Thus, dietary serine restriction might inhibit pancreatic carcinogenesis as well.

In summary, we demonstrate for the first time that metabolic reprogramming is an essential feature of acinar-to-ductal metaplasia the first step in pancreatic carcinogenesis. This reprogramming comprises increased glycolysis and a functional electron transport chain (ETC) to ensure serine synthesis and GSH synthesis. The observed changes are conserved among oncogene- and ligand-induced ADM formation. While the metabolic switch already takes place during the early stages of PDAC development, the diversification into metabolic subtypes must occur at later stages.

## Author contributions

T.N., N.W., M.C., S.U., R.O., S.L., M.D., T.M., M.J., K.S., W.S., H.E., R.M.S.

Conceptualization, T.N., H.E. and R.M.S.; Formal Analysis, T.N., S.U., R.O., T.M., W.S. and H.E.; Investigation, T.N., N.W., M.C., S.U., R.O., K.S. and W.S.; Resources, M.D., T.M. and M.J.; Writing – Original Draft, T.N., H.E. and R.M.S.; Visualization, T.N., K.S. and H.E.; Supervision, M.T., K.S., H.E., and R.M.S.; Project Administration, H.E. and R.M.S.; Funding Acquisition, T.M., K.S., R.M.S.

## Acknowledgements

We would like to thank Katrin Böttcher, Maximilian Reichert and Michael Quante for fruitful discussions and critical reading of the manuscript. T.N., N.W. K.S., H.E. and R.M.S. were supported by Deutsche Forschungsgemeinschaft (DFG)-SFB1321. T.M. was supported by Austrian Science Fund (FWF) grants P28854, I3792, DK-MCD W1226, DOC-130; Austrian Research Promotion Agency (FFG) Grants 864690 and 870454; the Integrative Metabolism Research Center Graz; Austrian Infrastructure Program 2016/2017, the Styrian Government (Zukunftsfonds, doc.funds program), the City of Graz and BioTechMed-Graz (Flagship project DYNIMO). S.U. was trained within the frame of the PhD program Metabolic and Cardiovascular Disease (DK-MCD).

## Declaration of interests

None.

## Material and Methods

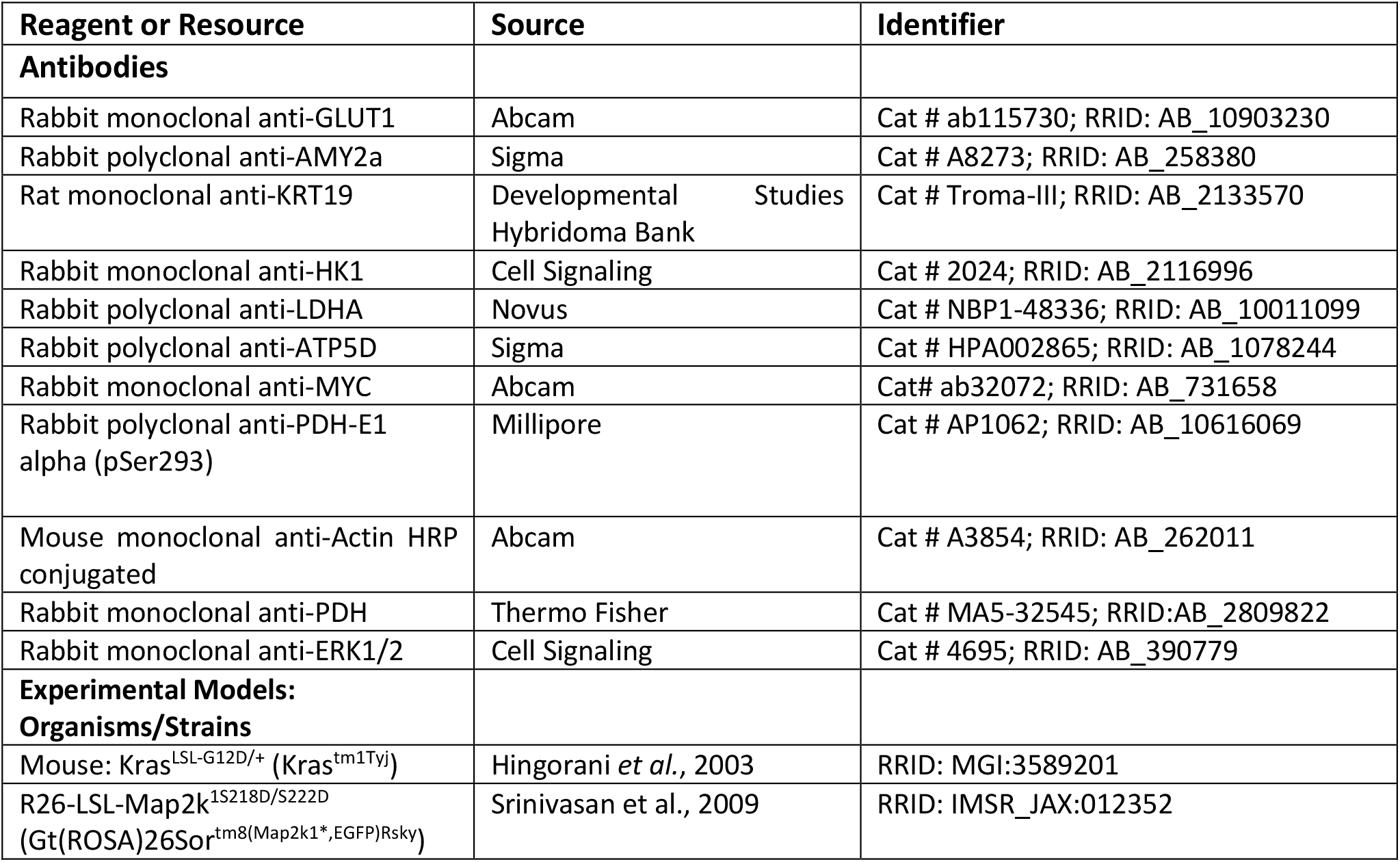
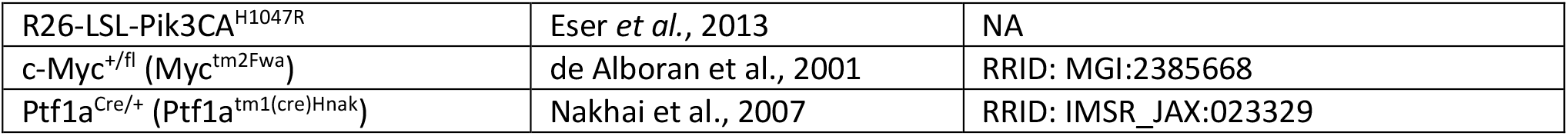

### Mouse strains, ethics and housing conditions

Kras^LSL-G12D/+^ (Kras^tm1Tyj^)^12^, R26-LSL-Map2k1^S218D/S222D^ (Gt(ROSA)26Sor^tm8(Map2k1*,EGFP)Rsky^)^40^, R26-LSL-Pik3CA^H1047R 13^ and c-Myc^+/fl^ (Myc^tm2Fwa^) ^41^ have been described before. These strains were interbred with Ptf1a^Cre/+^ (Ptf1a^tm1(cre)Hnak^) ^42^ mice to obtain mice with pancreas-specific alteration of selected pathways: Ptf1a^Cre/+^; Kras^LSL-G12D/+^ (Kras^G12D^), Ptf1a^Cre/+^; R26-LSL-Pik3CA^H1047R^ (Pi3k^CAH1047R^), Ptf1a^Cre/+^; R26-LSL-Map2k1^S218D/S222D^ (Mek1dd), Ptf1a^Cre/+^; c-Myc^+/fl^ (Myc^+/-^). All mouse strains were maintained in a mixed genetic background (C57BL6/ FVB/129) and housed in an SPF facility with a 12h:12h light:dark cycle and ad libitum access to food and water. Both sexes were included in the analyses. For *in vitro* ADM experiments, mice of 3-6 weeks of age were used. For *in vivo* ADM experiments, mice of 10 weeks of age were used. All animal studies were conducted in compliance with European guidelines for the care and use of laboratory animals and were approved by the Regierung von Oberbayern.

### Murine acinar cell isolation

Murine pancreata were harvested, cut into pieces and digested with SOL2 (1.2 mg/ml Collagenase from *Clostridium histolyticum* (C2139, Sigma) in McCoy’s 5A Medium, 0.02% Trypsin inhibitor from soy bean, 0.1% BSA (all Sigma)) twice for 10 min. Cells were passed through a 100 μm mesh and washed with SOL1 (McCoy’2 5A Medium, 0.02% Trypsin inhibitor from soy bean, 0.1% BSA) and spun down at 14 xg. Afterwards, cells were recovered in culture medium (Waymouth’s MB752/1 medium + 0.1% FCS, 1x Insulin-Transferrin-Selenium, 50 μg/mL Bovine Pituitary Extract, 10 mM HEPES (all Gibco), 0.1% BSA, 0.01% Trypsin inhibitor from soy bean, 2.6 mg/ml NaHCO^3^ (Merck) + 30% FCS and incubated for 30–60 min at 37°C. The endpoint of the incubation was defined as timepoint ‘d0’.

### Human acinar cell isolation

Healthy human pancreata were obtained from Surgery Department of the MRI. After soaking in SOL1 (McCoy’2 5A Medium, 0.02% Trypsin inhibitor from soy bean, 0.1% BSA) for 10 min, RT, specimens were cut into pieces and digested twice with 10 ml/g tissue of 50 U/ml collagenase (CLSPA Worthington Biochemical Corporation) in SOL1 for 10 min, stirred at 37°C. Cells were passed through a 200 μm cell strainer, washed with SOL1 and centrifuged at 157 xg. Cells were recovered in culture medium (see murine acinar cells) + 30% FCS and incubated for 30-60 min at 37°C. Afterwards, cells were collected at 14 xg for collagen embedding.

### *In vitro* transdifferentiation assay

Isolated acinar cells were recovered in culture medium, mixed at a ratio of 1:1 with neutralized collagen I from rat tail (Corning) and plated on a solid layer of 2.5 mg/ml collagen containing 10% 10x PBS. After solidification, an additional layer of collagen was added. Culture medium was added on top of the three-layer culture system. On day 1, the medium was changed. Transdifferentiation rates were determined at d2 (Mek1dd, Pi3k^CAH1047R^) or d3 (Kras^G12D^, stimulated wild-type). To determine the transdifferentiation rate all grape-like structures were counted as acini; hollow, spherical structures were counted as ADM structures. The transdifferentiation rate were calculated as the ratio of ADM structures to the total amount of structures (acini and ADM structures). Stimuli and inhibitors were added during the whole time of cultivation: 100 ng/ml TGFα (R&DSystems); 20 ng/ml IL17a (PeproTech); 30, 60, 120, 240 μM WZB117; 10 nM Oligomycin; 500 nM BPTES (all Selleckchem); 100 nM Antimycin A; 10 μM Rotenone; 2, 4, 8 mM 2-d-deoxyglucose; 200, 300, 400 μM CPI-613; 35 nM CCCP; 25, 50, 100 μM 10058-F4; 0.4 mM serine; 1 mM nicotinamide (all Sigma); 100, 200 μM Galloflavin; 10 μM NTC-502 (Tocris); 100 μM buthionine sulfoximine (Enzo).

### Cytotoxicity assay

To determine treatment induced cytotoxicity, LDH release into the medium was measured. All supernatants from the transdifferentiation assay were collected. Acinar cells lysed with 2% TritonX-100 (Sigma) were used as positive control; supernatant from medium-treated acinar cells as negative control. LDH activity was assayed using the Cytotoxicity Detection Kit (Roche) according to the manufacturer’s protocol.

### OCR and ECAR measurement

Seahorse measurements were performed as described before^43^. Briefly, acinar explants were seeded in collagen one day before measurement. 1 h before the measurement, the medium was changed to XF assay medium (Agilent) containing 5 g/l glucose. One assay cycle consisted of 1 min mixing, 2 min waiting and 4 min measuring. After basal respiration measurements (5 cycles), oligomycin was added (2 μg/ml) to inhibit ATP production and proton leak respiration measured (4 assay cycles). Afterwards, carbonyl cyanide m-chlorophenylhydrazone (CCCP) (1 μM) and pyruvate (5 mM) were added to determine the maximal respiratiory rate (4 cycles). Addition of rotenone (2.5 μM) and antimycin A (2.5 μM) served to fully inhibit the respiratory chain (3 cycles). 2-deoxyglucose (2DG) (100 mM) was added to inhibit glycolysis and correct for non-glycolytic acidification (6 cycles). ATP production rates were calculated as published previously^44^.

### Immunofluorescence

Methanol-fixed acinar cells in collagen gels were washed with PBS + 0.5% TritonX-100 (PBST) and blocked with 5% BSA in PBST for 1 h. Blocked cells were stained with antibodies against AMY2A (1:300, Sigma, A8273) or GLUT1 (1:300, abcam, ab115730) and KRT19 (1:500, DSHB, TROMA-III) overnight, consecutively. Secondary antibodies (Alexa488, A11006 and Alexa568, A11036 both Invitrogen) were incubated overnight. Gels were mounted with HardSet with DAPI (H-1500, Vectashield).

### Immunohistochemistry

Evaluation of expression of Hexokinase1, GLUT1 and LDHA was evaluated on human tissues using tissue microarrays (TMAs). For TMA construction, tissue blocks from in total 63 patients from the archive of the Institute of Pathology of TUM were selected, containing either PanIN and/or acinar-to-ductal metaplasia. The slides were annotated and TMAs were constructed using a tissue microarrayer (Beecher Instruments). For murine tissues, additionally expression of ATP5D was evaluated immunohistochemically. 2 μm thin slides were then prepared and immunohistochemistry was performed on a Bond Rxm system (Leica) with primary antibodies against Hexokinase 1 (1:500, Cell signaling, 2024), LDHA (1:10.000, Novus Biologicals, NBP1-48336), GLUT1 (1:750, abcam, ab115730) or ATP5D (1:100, Sigma Aldrich, HPA002865). Briefly, after deparaffinization, slides were pretreated with antigen retrieval solution 1 (ER1, Leica) for 30 or 40 minutes and primary antibody binding was visualized using a Polymer Refine Detection Kit (Leica). Slides were scanned with an AT2 (Leica) scanning system and evaluated semiquantitatively by a board-certified pathologist (KS). Marker expression in human tissue was scored with respect to the specific pathology using a 4-tier scoring system (0-no expression, 1-slight expression, 2-moderate expression, 3-strong expression). For the murine specimen, expression for all marker was evaluated with respect to normal acinar tissue in comparison to acinar-to-ductal metaplasia (ADM) using a 6-tier scoring system reflecting expression frequency and intensity (from 0-no expression to 5-strong expression in all cells).

### Isolation of RNA

Cultivated acinar cells were explanted from the collagen gel using SOL2 (see murine acinar isolation). Afterwards, cells were pelleted at 300 g and lysed in 200 μL homogenisation buffer including 2% 1-thioglycerol. The isolation was done using the Maxwell Kit (all Promega) according to the manufacture’s protocol.

### RNASeq

Library preparation for bulk-sequencing of poly(A)-RNA was done as described previously^45^. Briefly, barcoded cDNA was generated with a Maxima RT polymerase (ThermoFisher) using oligo-dT primer containing barcodes, unique molecular identifiers (UMIs) and an adaptor. Ends of the cDNAs were extended by a template switch oligo (TSO) and full-length cDNA was amplified with primers binding to the TSO-site and the adaptor. NEB UltraII FS kit was used to fragment cDNA. After end repair and A-tailing, a TruSeq adapter was ligated and 3’-end-fragments were amplified using primers with Illumina P5 and P7 overhangs. In comparison to Parekh et al. (2016), the P5 and P7 sites were exchanged to allow sequencing of the cDNA in read1 and barcodes and UMIs in read2. The library was sequenced on a NextSeq 500 (Illumina) with 67 cycles for the cDNA in read1 and 16 cycles for the barcodes and UMIs in read2. Data was processed using the published Drop-seq pipeline (v1.0) to generate sample- and gene-wise UMI tables^46^. Reference genome (GRCm38) was used for alignment. Transcript and gene definitions were used according to the GENCODE Version M25. RNASeq data were normalized with the Deseq2 package in R.

### Western Blot

Acinar cells were lysed with RIPA buffer containing protease and phosphatase inhibitors. Protein extracts were separate using SDS-PAGE, transferred to a nitrocellulose membrane and tested with the following antibodies: GLUT1 (ab115730), c-MYC (ab32072) (both abcam), LDHA (NBP1-48336, Novobio), AMY2a (A8273), β-ACTIN (#A3854) (all sigma), p-PDH (AP1062, Millipore), KRT19 (Troma III, DSHB), PDH (MA5-32545, Invitrogen), ERK1/2 (#4695, CellSignaling), anti-rabbit-HRP (GE_NA934V) and anti-rat-HRP (GE_NA935V) (both GE Healthcare).

### Metabolome analysis by NMR

For extraction of metabolites from cells and tissue, samples were homogenized in 600 μL ice-cold 66% MetOH in H_2_O using a Precellys24 tissue homogenizer and incubated for 1 h at - 20°C. For metabolite extraction from medium, 200 μl of medium were mixed with 400 μl ice-cold MetOH and incubated for 1 h at -20°C. Afterwards, all samples were centrifuged at 17949 g for 30 min at 4°C and supernatants were lyophilized at <1 Torr, 213 g, 25°C for 10 h. For the NMR experiments, extracts were dissolved in 500 μl NMR buffer (0.08 M Na_2_HPO_4_, 5 mM TSP (3-(trimethylsilyl) propionic acid-2,2,3,3-d_4_ sodium salt), 0.04 % (w/v) NaN_3_ in D_2_O, pH 7.4) and transferred to 5 mm NMR tubes. The analysis was run on a 600 MHz Bruker Avance Neo NMR spectrometer at 310 K. _1_H-1D-NMR spectra were acquired with a pre-saturation for water suppression (cpmgpr1d, 512 scans, 73728 points in F1, 12019.230 Hz spectral width, 1024 transients, recycle delay 4s)^47^. NMR spectral data were processed as previously described^48^. Spectra analysis was performed using Brucker Topspin, Matlab and MetaboAnalyst^49^. For metabolite identification, Chenomx NMR Suite 8.4 was used.

### Mass spectrometry and isotope tracing

Isolated acinar were incubated with either complete medium (including designated inhibitors) or glucose-free medium (ThermoFisher) with 27.7 mM [^13^C_6_]-glucose (Sigma). Cells were labeled from 0-2 h or 22-24 h after isolation. Labelled cells were washed with cold PBS and pellets were frozen at -80°C.

Water-soluble metabolites were extracted with 0.5 ml ice-cold MeOH/H_2_O (80/20, v/v) containing 0.01 μM lamivudine (Sigma-Aldrich) and 0.5 μM sucrose (Sigma-Aldrich). After centrifugation of the homogenates, supernatants were transferred to a RP18 SPE column (Phenomenex) that had been activated with 0.5 ml CH_3_CN and conditioned with 0.5 mL of MeOH/H_2_O (80/20, v/v). The eluate was dried in a centrifugal evaporator (Savant) and dissolved in 50 μL 5 mM NH_4_OAc in CH_3_CN/H_2_O (50/50, v/v).

Metabolites were analysed by LC-MS using the following settings:

For LC-MS analysis 3 μl of sample was applied to a ZIC-cHILIC column (SeQuant ZIC-cHILIC, 3 μm, 100*2.1 mm). Metabolites were separated with Solvent A, consisting of 5 mM NH_4_OAc in CH_3_CN/H_2_O (5/95, v/v) and solvent B consisting of 5 mM NH_4_OAc in CH_3_CN/H_2_O (95/5, v/v) at a flow rate of 200 μl/min at 30°C by LC using a DIONEX Ultimate 3000 UPLC system (Thermo Scientific), applying a linear gradient starting after 2 min with 100% solvent B decreasing to 40% solvent B over 23 min, followed by 17 min 40% solvent B and a linear increase to 100% solvent B over 1 min Recalibration of the column was achieved by 7 min prerun with 100% solvent B.

All MS-analyses were performed on a high-resolution QExactive mass spectrometer (ThermoScientific) in alternating positive- and negative full MS mode with a Scan Range of 69.0 - 1000 m/z at 70K Resolution and the following ESI source parameters: Sheath gas: 30, auxiliary gas: 10, sweep gas: 3, Aux Gas Heater temperature: 120 °C. Spray voltage: 2.5 kV in positive ion mode and 3.6 kV in negative ion mode, Capillary temperature: 320 °C, S-lens RF level: 55.0. Signal determination and quantitation was performed using TraceFinder™ Software Version 3.3 (ThermoFisher). All analyses were performed with five independent biological replicates.

### Statistical analysis

The graphical arrangement of data and statistically testing for significance was done using GraphPad Prism 8 (GraphPad Software). For parametric data a Student’s t-test or an ANOVA was performed. For non-parametric data (histological scores) a Kruskal-Wallis test was used.

## Figure Legends

**Sup Fig. 1.**
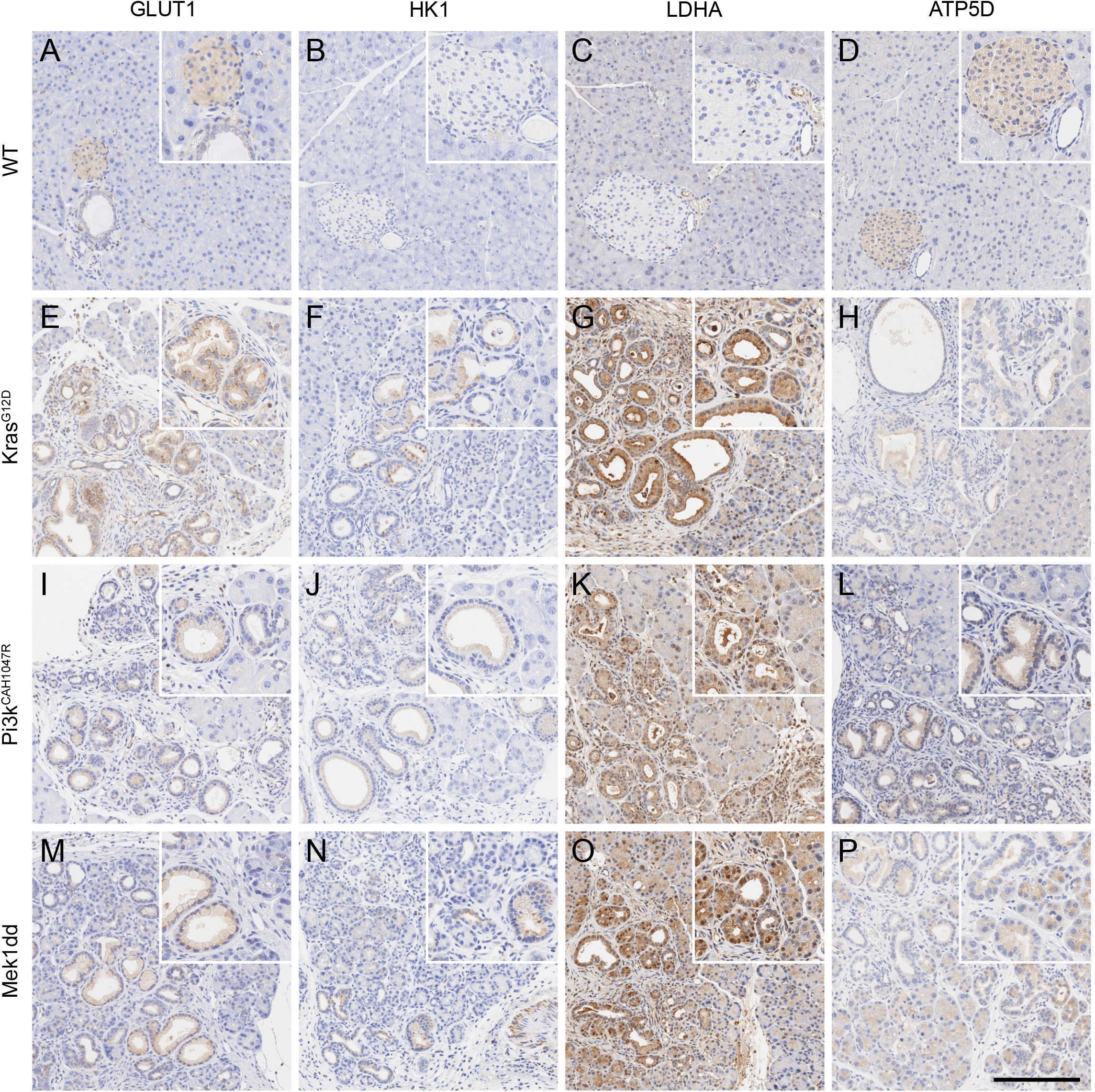
Glycolysis is upregulated during ADM *in vivo*. **A-R**. Immunohistochemistry for GLUT1 (**A, E, I, M**), HK1 (**B, F, J, N**), LDHA (**C, G, K, O**) and ATP5D (**D, H, L, P**) on pancreatic tissue from 10-week-old mice. Scale bar = 200 μm.

**Sup Fig. 2.**
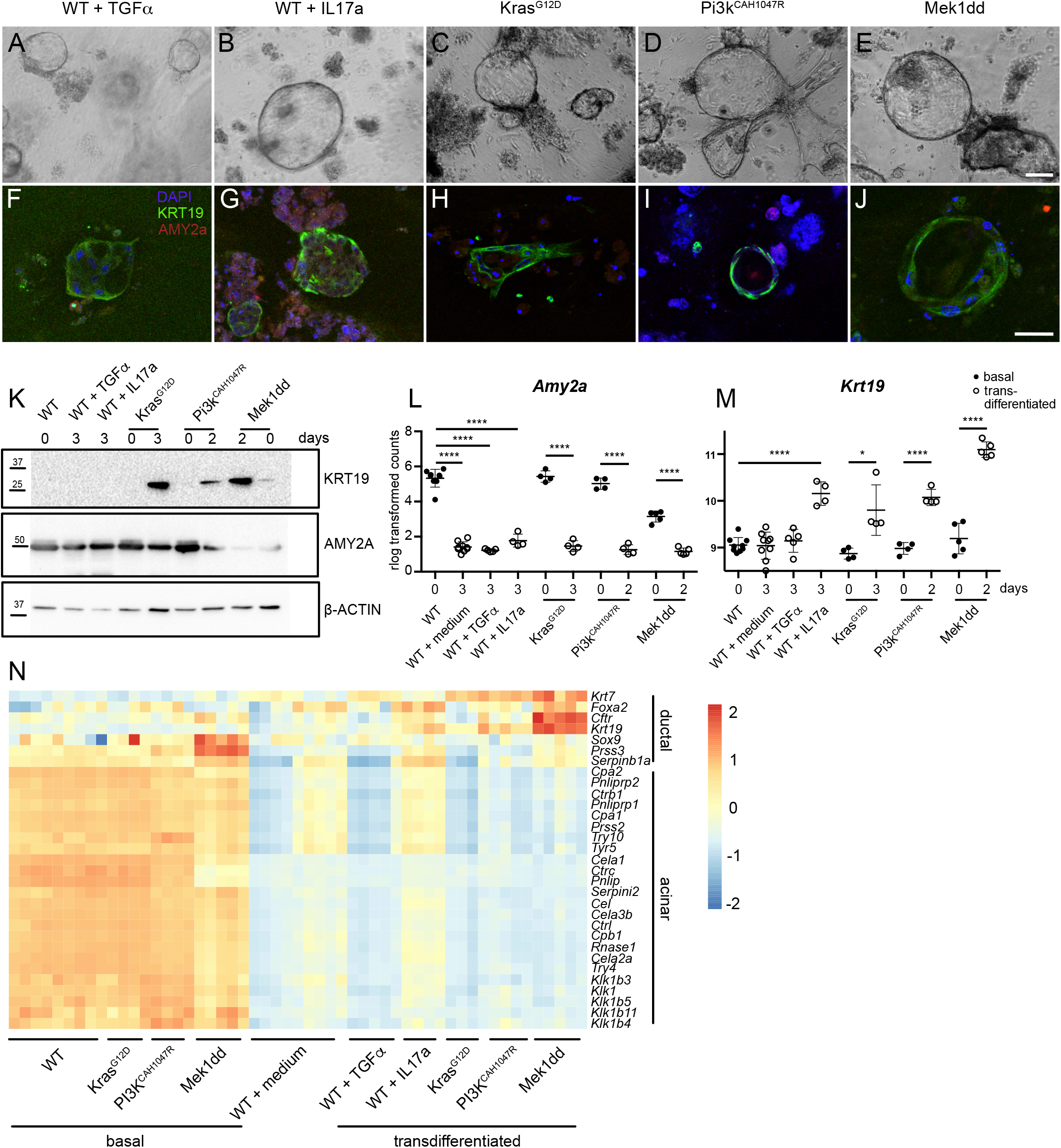
Different stimuli show comparable ADM formation *in vitro*. **A-E**. Light microscopical images of transdifferentiated acinar cells after collagen culture. Scale bar = 100 μm. **F-J**. Immunofluorescence images of transdifferentiated acinar cells stained for KRT19 (green), AMY2A (red) and DAPI (blue). Scale bar = 50 μm. **K**. Protein expression of KRT19 and AMY2A in either freshly isolated or transdifferentiated acinar cells. b-ACTIN served as a loading control. **L**. Expression of Amy2a in freshly isolated or cultured acinar cells. **M**. Expression of Krt19 in freshly isolated or cultured acinar cells. **N**. RNA expression data for acinar or ductal specific transcripts in freshly isolated (“basal”) or cultured acinar cells (“transdifferentiated”). Results are mean +/- SD, *p<0.05, **p<0.01, ***p<0.001, ****p<0.0001.

**Sup Fig. 3.**
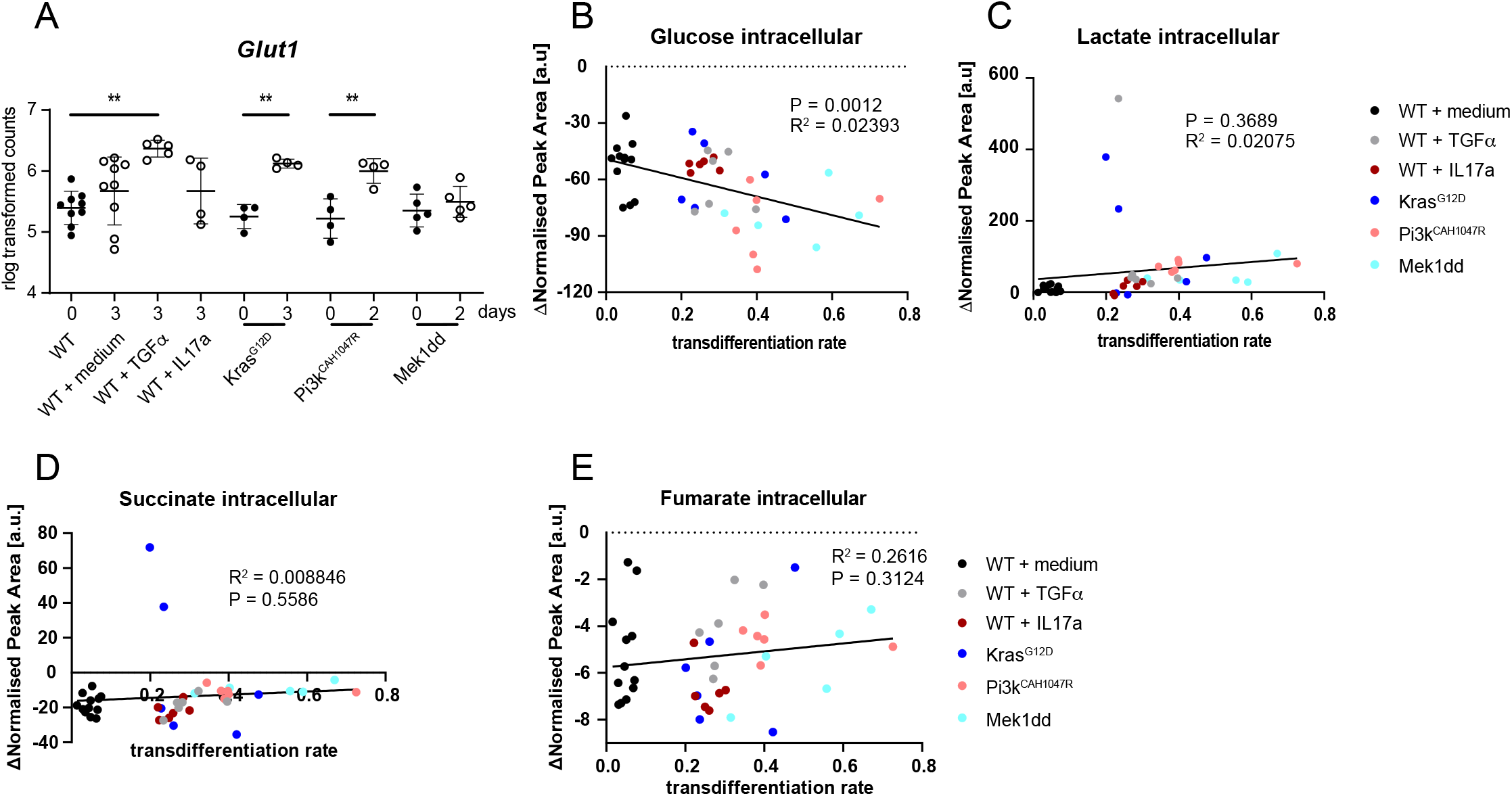
Downregulation of OxPhos and uptake of amino acids during ADM formation. **A**. RNA expression of the glucose transporter 1 (*Glut1*) in either freshly isolated or transdifferentiated acinar cells. **B-H**. Amounts of glucose (**B**), lactate (**C**) and succinate (**D**) in transdifferentiated acinar cells. Values of freshly isolated acinar cells were subtracted from the respective samples.

**Sup Fig. 4.**
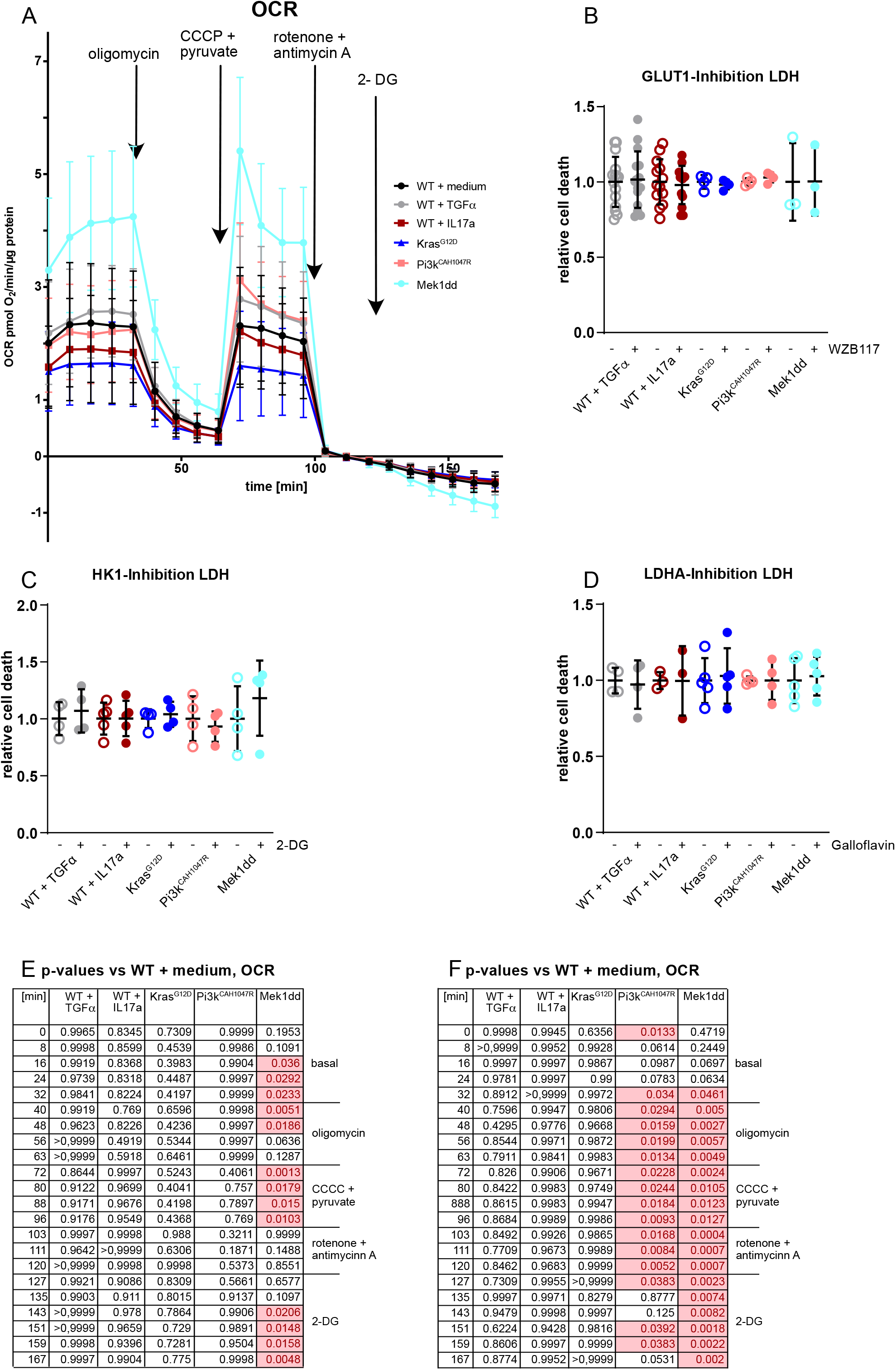
Seahorse measurements and LDH release assay. **A**. Oxygen consumption rate of acinar cells 24 h after isolation. Cells were treated with the displayed inhibitors at the given timepoints. All values were normalised by the protein content per well (N = 7-9). **B, C, D**. LDH release from treated acinar cells as indicator of cytotoxicity. Values for the untreated controls and the highest concentrations of WZB117 (**B**), 2-deoxyglucose (2-DG) (**C**) and Galloflavin (**D**), respectively. **E**,**F** p values for OCR (**E**) and ECAR (**F**) per time point. Values were compared to WT + medium. All values have been normalised to the untreated controls. Results are mean +/- SD.

**Sup Fig. 5.**
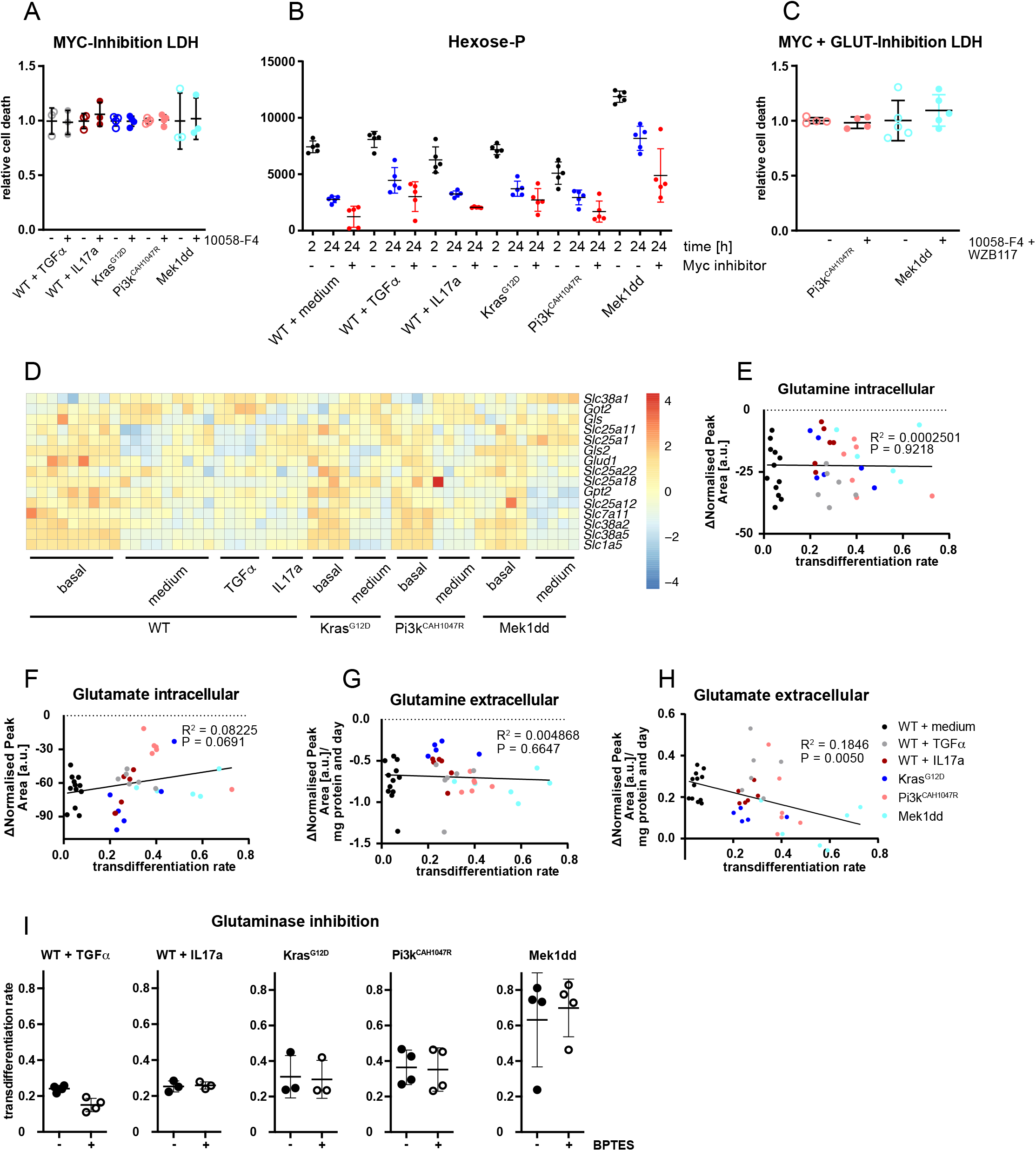
Glutaminolysis is dispensable for ADM. **A**. LDH release from treated acinar cells as indicator of cytotoxicity. Values for the untreated controls and treatment with 10058-F4. All values have been normalized to the untreated controls. **B**. LDH release from treated acinar cells as indicator of cytotoxicity in untreated controls and after treatment with the combination of 10058-F4 and WZB117. All values have been normalised to the untreated controls. **C**. Total amounts of hexose-P in acinar cells after treatment with 50 μM 10058-F4. **D**. RNA expression data for glutaminolysis associated genes (transporters and enzymes) in freshly isolated (“basal”) and cultured acinar cells (“transdifferentiated”). **E, F**. Amounts of glutamine (**E**) and glutamate (**F**) in transdifferentiated acinar cells and their supernatants. Values of freshly isolated acinar cells were subtracted from the respective samples. G, **H**. Amounts of glutamine (**G**) and glutamate (**H**) in the supernatants of transdifferentiated acinar cells. Values of unused medium were subtracted from the respective samples. Medium values were normalised by the pellet weight of the acinar cells. **I**. Transdifferentiation rates after treatment with the glutaminase inhibitor BPTES. Results are mean +/- SD, *p<0.05.

**Sup Fig. 6.**
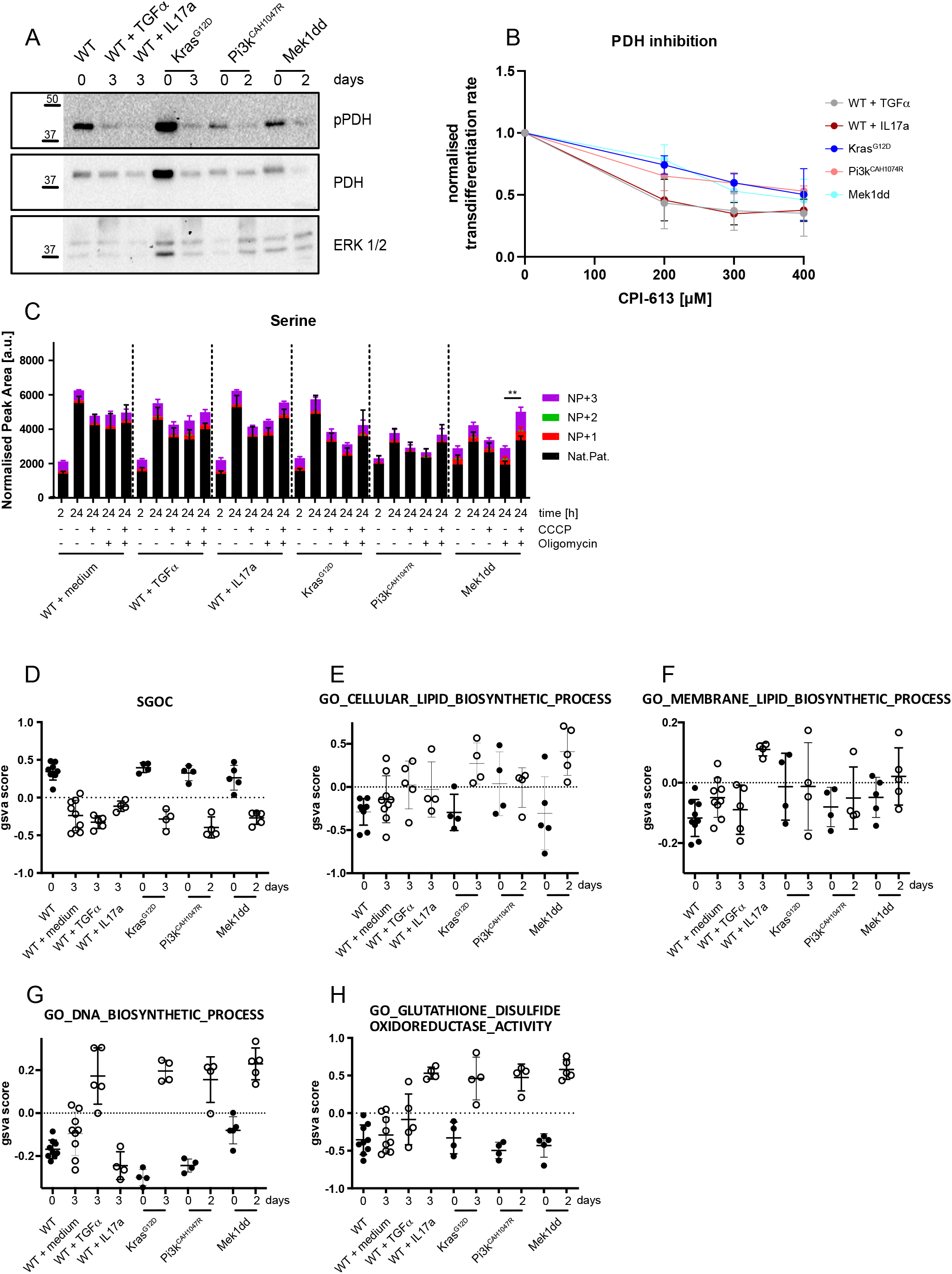
Mitochondrial regulator PDH is activated and required for ADM and serine levels correlate with mitochondrial activity. **A**. Western Blot for (p-)PDH of either freshly isolated or transdifferentiated acinar cells. ERK1/2 served as a loading control. **B**. Transdifferentiation rates after treatment with the PDH inhibitor CPI-613 (N = 3-11). **C**. Isotopologue distribution of intracellular serine amounts in acinar cells 2 or 24 h after isolation. Acinar cells were treated either with oligomycin or CCCP or the combination of both (N = 5). **D-H**. Enrichment scores for the genesets SGOC (Serine-Gylcin-One-Carbon) (**D**), GO_CELLULAR_LIPID_BIOSYNTHETIC_PROCESS (**E**), GO_MEMBRANE_LIPID_BIOSYNTHETIC_PROCESS (**F**), GO_DNA_BIOSYNTHETIC_PROCESS (**G**) and GO_GLUTATHIONE_DISULFIDE_OXIDOREDUCTASE_ACTIVITY (**H**) in RNASeq of freshly isolated and cultured acinar cells. NP = normal pattern. Results are mean +/- SD, **p<0.01.

## Notes

### Competing Interest Statement

The authors have declared no competing interest.

### Summary of Updates

Added substantial data concerning serine importance

## References

1 Hanahan, D. & Weinberg, R. A. The hallmarks of cancer. Cell 100, 57–70 (2000). https://doi.org:10.1016/s0092-8674(00)81683-9

2 Warburg, O., Wind, F. & Negelein, E. The Metabolism of Tumors in the Body. J Gen Physiol 8, 519–530 (1927). https://doi.org:10.1085/jgp.8.6.519

3 Ying, H. et al. Oncogenic Kras maintains pancreatic tumors through regulation of anabolic glucose metabolism. Cell 149, 656–670 (2012). https://doi.org:10.1016/j.cell.2012.01.058

4 Viale, A. et al. Oncogene ablation-resistant pancreatic cancer cells depend on mitochondrial function. Nature 514, 628–632 (2014). https://doi.org:10.1038/nature13611

5 Valle, S. et al. Exploiting oxidative phosphorylation to promote the stem and immunoevasive properties of pancreatic cancer stem cells. Nat Commun 11, 5265 (2020). https://doi.org:10.1038/s41467-020-18954-z

6 Hezel, A. F., Kimmelman, A. C., Stanger, B. Z., Bardeesy, N. & Depinho, R. A. Genetics and biology of pancreatic ductal adenocarcinoma. Genes Dev 20, 1218–1249 (2006). https://doi.org:10.1101/gad.1415606

7 Wagner, M., Luhrs, H., Kloppel, G., Adler, G. & Schmid, R. Malignant transformation of duct-like cells originating from acini in transforming growth factor α transgenic mice. GASTROENTEROLOGY 115, 1254–1262 (1998). https://doi.org:10.1016/s0016-5085(98)70098-8

8 Zhu, L., Shi, G., Schmidt, C. M., Hruban, R. H. & Konieczny, S. F. Acinar cells contribute to the molecular heterogeneity of pancreatic intraepithelial neoplasia. Am J Pathol 171, 263–273 (2007). https://doi.org:papers2://publication/doi/10.2353/ajpath.2007.061176

9 Kopp, J. L. et al. Identification of Sox9-dependent acinar-to-ductal reprogramming as the principal mechanism for initiation of pancreatic ductal adenocarcinoma. Cancer Cell 22, 737–750 (2012). https://doi.org:10.1016/j.ccr.2012.10.025

10 Huang, L. et al. Commitment and oncogene-induced plasticity of human stem cell-derived pancreatic acinar and ductal organoids. Cell Stem Cell 28, 1090-1104.e1096 (2021). https://doi.org:10.1016/j.stem.2021.03.022

11 Dai, S. et al. Glycolysis promotes the progression of pancreatic cancer and reduces cancer cell sensitivity to gemcitabine. Biomed Pharmacother 121, 109521 (2020). https://doi.org:10.1016/j.biopha.2019.109521

12 Hingorani, S. R. et al. Preinvasive and invasive ductal pancreatic cancer and its early detection in the mouse. Cancer Cell 4, 437–450 (2003). https://doi.org:papers2://publication/uuid/C4F63D0D-F712-4563-913C-FF456289C45E

13 Eser, S. et al. Selective Requirement of PI3K/PDK1 Signaling for Kras Oncogene-Driven Pancreatic Cell Plasticity and Cancer. Cancer Cell (2013). https://doi.org:papers2://publication/doi/10.1016/j.ccr.2013.01.023

14 McAllister, F. et al. Oncogenic Kras activates a hematopoietic-to-epithelial IL-17 signaling axis in preinvasive pancreatic neoplasia. Cancer Cell 25, 621–637 (2014). https://doi.org:10.1016/j.ccr.2014.03.014

15 Murtaugh, L. C. & Keefe, M. D. Regeneration and repair of the exocrine pancreas. Annu Rev Physiol 77, 229–249 (2015). https://doi.org:10.1146/annurev-physiol-021014-071727

16 Dejure, F. R. & Eilers, M. MYC and tumor metabolism: chicken and egg. EMBO J 36, 3409–3420 (2017). https://doi.org:10.15252/embj.201796438

17 Mehrmohamadi, M., Liu, X., Shestov, A. A. & Locasale, J. W. Characterization of the usage of the serine metabolic network in human cancer. Cell Rep 9, 1507–1519 (2014). https://doi.org:10.1016/j.celrep.2014.10.026

18 Pavlova, N. N. & Thompson, C. B. The Emerging Hallmarks of Cancer Metabolism. Cell Metab 23, 27–47 (2016). https://doi.org:10.1016/j.cmet.2015.12.006

19 Pinho, A. V. et al. Sirtuin 1 stimulates the proliferation and the expression of glycolysis genes in pancreatic neoplastic lesions. Oncotarget 7, 74768–74778 (2016). https://doi.org:10.18632/oncotarget.11013

20 Rong, Y. et al. Lactate dehydrogenase A is overexpressed in pancreatic cancer and promotes the growth of pancreatic cancer cells. Tumour Biol 34, 1523–1530 (2013). https://doi.org:10.1007/s13277-013-0679-1

21 Voronina, S. G. et al. Dynamic changes in cytosolic and mitochondrial ATP levels in pancreatic acinar cells. GASTROENTEROLOGY 138, 1976–1987 (2010). https://doi.org:10.1053/j.gastro.2010.01.037

22 Bauduin, H., Colin, M. & Dumont, J. E. Energy sources for protein synthesis and enzymatic secretion in rat pancreas in vitro. Biochim Biophys Acta 174, 722–733 (1969). https://doi.org:10.1016/0005-2787(69)90301-3

23 Halangk, W. et al. Effect of supramaximal cerulein stimulation on mitochondrial energy metabolism in rat pancreas. Pancreas 16, 88–95 (1998). https://doi.org:10.1097/00006676-199801000-00014

24 Mayer, D., Klimek, F., Rempel, A. & Bannasch, P. Hexokinase expression in liver preneoplasia and neoplasia. Biochem Soc Trans 25, 122–127 (1997). https://doi.org:10.1042/bst0250122

25 Ahn, Y. S., Zerban, H. & Bannasch, P. Expression of glucose transporter isoforms (GLUT1, GLUT2) and activities of hexokinase, pyruvate kinase, and malic enzyme in preneoplastic and neoplastic rat renal basophilic cell lesions. Virchows Arch B Cell Pathol Incl Mol Pathol 63, 351–357 (1993). https://doi.org:10.1007/BF02899283

26 Sakashita, M. et al. Glut1 expression in T1 and T2 stage colorectal carcinomas: its relationship to clinicopathological features. Eur J Cancer 37, 204–209 (2001). https://doi.org:10.1016/s0959-8049(00)00371-3

27 Nie, M. et al. Evolutionary metabolic landscape from preneoplasia to invasive lung adenocarcinoma. Nat Commun 12 (2021). https://doi.org:10.1038/s41467-021-26685-y

28 Zhang, F. et al. Growing Human Hepatocellular Tumors Undergo a Global Metabolic Reprogramming. Cancers (Basel) 13 (2021). https://doi.org:10.3390/cancers13081980

29 Son, J. et al. Glutamine supports pancreatic cancer growth through a KRAS-regulated metabolic pathway. Nature 496, 101–105 (2013). https://doi.org:10.1038/nature12040

30 Bhutia, Y. D. & Ganapathy, V. Glutamine transporters in mammalian cells and their functions in physiology and cancer. Biochim Biophys Acta 1863, 2531–2539 (2016). https://doi.org:10.1016/j.bbamcr.2015.12.017

31 Walz, S. et al. Activation and repression by oncogenic MYC shape tumour-specific gene expression profiles. Nature 511, 483–487 (2014). https://doi.org:10.1038/nature13473

32 Shroff, E. H. et al. MYC oncogene overexpression drives renal cell carcinoma in a mouse model through glutamine metabolism. Proc Natl Acad Sci U S A 112, 6539–6544 (2015). https://doi.org:10.1073/pnas.1507228112

33 Dey, P. et al. Oncogenic KRAS-Driven Metabolic Reprogramming in Pancreatic Cancer Cells Utilizes Cytokines from the Tumor Microenvironment. Cancer Discov 10, 608–625 (2020). https://doi.org:10.1158/2159-8290.CD-19-0297

34 Sancho, P. et al. MYC/PGC-1alpha Balance Determines the Metabolic Phenotype and Plasticity of Pancreatic Cancer Stem Cells. Cell Metab 22, 590–605 (2015). https://doi.org:10.1016/j.cmet.2015.08.015

35 Yang, M. & Vousden, K. H. Serine and one-carbon metabolism in cancer. Nat Rev Cancer 16, 650–662 (2016). https://doi.org:10.1038/nrc.2016.81

36 Yang, L. et al. Serine Catabolism Feeds NADH when Respiration Is Impaired. Cell Metab 31, 809–821 e806 (2020). https://doi.org:10.1016/j.cmet.2020.02.017

37 Diehl, F. F., Lewis, C. A., Fiske, B. P. & Vander Heiden, M. G. Cellular redox state constrains serine synthesis and nucleotide production to impact cell proliferation. Nat Metab 1, 861–867 (2019). https://doi.org:10.1038/s42255-019-0108-x

38 Banh, R. S. et al. Neurons Release Serine to Support mRNA Translation in Pancreatic Cancer. Cell 183, 1202-1218.e1225 (2020). https://doi.org:10.1016/j.cell.2020.10.016

39 Tajan, M. et al. Serine synthesis pathway inhibition cooperates with dietary serine and glycine limitation for cancer therapy. Nat Commun 12 (2021). https://doi.org:10.1038/s41467-020-20223-y

40 Srinivasan, L. et al. PI3 kinase signals BCR-dependent mature B cell survival. Cell 139, 573–586 (2009). https://doi.org:papers2://publication/doi/10.1016/j.cell.2009.08.041

41 de Alboran, I. M. et al. Analysis of C-MYC function in normal cells via conditional gene-targeted mutation. Immunity 14, 45–55 (2001). https://doi.org:10.1016/s1074-7613(01)00088-7

42 Nakhai, H. et al. Ptf1a is essential for the differentiation of GABAergic and glycinergic amacrine cells and horizontal cells in the mouse retina. Development 134, 1151–1160 (2007). https://doi.org:papers2://publication/doi/10.1242/dev.02781

43 Keuper, M. et al. Spare mitochondrial respiratory capacity permits human adipocytes to maintain ATP homeostasis under hypoglycemic conditions. FASEB J 28, 761–770 (2014). https://doi.org:10.1096/fj.13-238725

44 Mookerjee, S. A., Gerencser, A. A., Nicholls, D. G. & Brand, M. D. Quantifying intracellular rates of glycolytic and oxidative ATP production and consumption using extracellular flux measurements. J Biol Chem 292, 7189–7207 (2017). https://doi.org:10.1074/jbc.M116.774471

45 Parekh, S., Ziegenhain, C., Vieth, B., Enard, W. & Hellmann, I. The impact of amplification on differential expression analyses by RNA-seq. Sci Rep 6, 25533 (2016). https://doi.org:10.1038/srep25533

46 Macosko, E. Z. et al. Highly Parallel Genome-wide Expression Profiling of Individual Cells Using Nanoliter Droplets. Cell 161, 1202–1214 (2015). https://doi.org:10.1016/j.cell.2015.05.002

47 Vignoli, A. et al. High-Throughput Metabolomics by 1D NMR. Angewandte Chemie International Edition 58, 968–994 (2019). https://doi.org:10.1002/anie.201804736

48 Streese, L. et al. Metabolic profiling links cardiovascular risk and vascular end organ damage. Atherosclerosis 331, 45–53 (2021). https://doi.org:10.1016/j.atherosclerosis.2021.07.005

49 Chong, J. et al. MetaboAnalyst 4.0: towards more transparent and integrative metabolomics analysis. Nucleic Acids Res. 46, W486–W494 (2018). https://doi.org:10.1093/nar/gky310

